# CRISPRi-based screen of Autism Spectrum Disorder risk genes in microglia uncovers roles of *ADNP* in microglia endocytosis and synaptic pruning

**DOI:** 10.1101/2024.06.01.596962

**Authors:** Olivia M Teter, Amanda McQuade, Venus Hagan, Weiwei Liang, Nina M Dräger, Sydney M Sattler, Brandon B Holmes, Vincent Cele Castillo, Vasileios Papakis, Kun Leng, Steven Boggess, Tomasz J Nowakowski, James Wells, Martin Kampmann

**Affiliations:** Institute for Neurodegenerative Diseases, University of California, San Francisco, San Francisco, CA, USA; UC Berkeley-UCSF Graduate Program in Bioengineering, University of California, San Francisco, San Francisco, CA, USA; Department of Neurology, University of California, San Francisco, CA, USA; Developmental and Stem Cell Biology Graduate Program, University of California, San Francisco, San Francisco, CA, USA; Biomedical Sciences Graduate Program, University of California, San Francisco, San Francisco, CA, USA; Medical Scientist Training Program, University of California, San Francisco, San Francisco, CA, USA; Department of Anatomy, University of California, San Francisco, San Francisco, CA 94158, USA; Department of Psychiatry and Behavioral Sciences, University of California, San Francisco, San Francisco, CA 94158, USA; Department of Neurological Surgery, University of California, San Francisco, San Francisco, CA 94158, USA; Weill Institute for Neurosciences, University of California, San Francisco, San Francisco, CA 94158, USA; Eli and Edythe Broad Center for Regeneration Medicine and Stem Cell Research, University of California, San Francisco, San Francisco, CA 94158, USA; Department of Pharmaceutical Chemistry, University of California, San Francisco, San Francisco, CA, USA; Department of Biochemistry and Biophysics, University of California, San Francisco, San Francisco, CA, USA

## Abstract

Autism Spectrum Disorders (ASD) are a set of neurodevelopmental disorders with complex biology. The identification of ASD risk genes from exome-wide association studies and de novo variation analyses has enabled mechanistic investigations into how ASD-risk genes alter development. Most functional genomics studies have focused on the role of these genes in neurons and neural progenitor cells. However, roles for ASD risk genes in other cell types are largely uncharacterized. There is evidence from postmortem tissue that microglia, the resident immune cells of the brain, appear activated in ASD. Here, we used CRISPRi-based functional genomics to systematically assess the impact of ASD risk gene knockdown on microglia activation and phagocytosis. We developed an iPSC-derived microglia-neuron coculture system and high-throughput flow cytometry readout for synaptic pruning to enable parallel CRISPRi-based screening of phagocytosis of beads, synaptosomes, and synaptic pruning. Our screen identified *ADNP*, a high-confidence ASD risk genes, as a modifier of microglial synaptic pruning. We found that microglia with ADNP loss have altered endocytic trafficking, remodeled proteomes, and increased motility in coculture.

## INTRODUCTION

Autism is a spectrum of neurodevelopmental disorders defined by core features, such as social and communication impairments and repetitive or restrictive behaviors. Although there is behavioral and etiological diversity – with both environmental and genetic factors contributing to Autism Spectrum Disorder (ASD) risk – there are shared patterns of atypical brain development including dysregulated synaptic density^1–3^ and activation of microglia in people with ASD ^4–8^. However, it is not known if the observed activation of microglia in ASD is in response to abnormal neuronal cues or an intrinsic state of the microglia. Microglia play traditional immune regulatory roles as well as specialized roles to support circuit development including clearance of apoptotic neural progenitors^9,10^, synaptic remodeling^11–13^, and regulation of neuronal activity^14,15^. It is not known if the observed activation of microglia in ASD is in response to abnormal neuronal cues or an intrinsic state of the microglia. Thus, there is a critical need to focus research into which of these microglial functions may be altered in ASD.

Given the complexity and multi-cellular coordination in these developmental processes, it is difficult to illuminate mechanisms underpinning how ASD risk factors give rise to atypical brain development. However, progress in genome sequencing and de novo mutation analyses have led to the identification of large effect size genes implicated in ASD risk^16^. Since the identification of these risk genes, CRISPR-based genetic screens in human iPSC-derived two-and three-dimensional neural cultures *in vitro*, as well as in mice have been conducted. These studies have focused on phenotypes in the neural lineage, including neurogenesis, neuronal migration and differentiation, and progenitor cell proliferation^17–22^. A subset of ASD risk genes are expressed, albeit not enriched, in microglia, and whether their disruption contributes to ASD biology is unknown. In this work, we explore how these risk genes act in microglia by using CRISPRi-based functional genomics. Many ASD risk genes are thought to be haploinsufficient^23–25^ due to the evidence of increased burden of heterozygous de novo loss of function mutations, including protein truncating variants and deleterious missense variants. Therefore, targeting of these genes by CRISPRi, which results in a partial reduction in gene expression, more closely models a heterozygous loss of function than complete gene knockout. We used CRIPSRi to assess whether knocking down ASD risk genes in microglia alters their transcriptomic activation state and synaptic remodeling.

SIngle cell sequencing revealed that knockdown of many ASD risk genes shifts the activation profile of iPSC-derived microglia (iTF-Microglia). To characterize the functional impact of ASD risk genes, we developed a flow cytometry-based readout for synaptic pruning, an important process in neurodevelopment, using an engineered human iPSC-derived microglia-neuron coculture system. The development of this high-throughput synaptic pruning assay enabled us to conduct three pooled CRISPRi-based screens in parallel, assessing phagocytosis across beads, synaptosomes, and synaptic pruning. In our work, we define synaptic as the observation of synaptic material from neurons being taken up by microglia; however, the mechanism by which synaptic material from neurons is taken up by microglia is not well defined and the nuances of this term are discussed in a recent review^26^. Our comprehensive approach led to the identification of a subset of ASD risk genes that alter phagocytosis across substrates. Surprisingly, we found one gene, *ADNP*, that specifically regulates synaptic pruning. Focused molecular and cellular characterization of iTF-Microglia with *ADNP* knockdown revealed that *ADNP* loss disrupts endocytic trafficking, remodels their surface proteome, and alters interactions with neurons. Through this focused investigation, we provide evidence that the disruption of ASD risk gene function in microglia can alter the microglial activation state and interactions with neurons.

## MATERIALS AND METHODS

### Human iPSC culture

iPSCs were cultured on Matrigel (Corning; Cat. No. 356231) coated vessels in StemFlex Media (Gibco; Cat. No. A3349401) until 80% confluent. Upon confluency, cells were washed with DPBS (Gibco; Cat. No. 14190-144), lifted using StemPro Accutase Cell Dissociation Reagent (GIBCO/Thermo Fisher Scientific; Cat. No. A11105-01) for 3 minutes at 37 °C, diluted with DPBS, and pelleted by 5 minutes of centrifugation at 220g. The supernatant was aspirated, cells counted, resuspended in StemFlex supplemented with 10 µM Y-27632 dihydrochloride ROCK inhibitor (Tocris; Cat. No. 1254), and plated on Matrigel coated vessels. StemFlex was replenished daily.

### Human iPSC-derived microglia (iTF-Microglia) with CRISPRi compatibility

iPSC-derived microglia (iTF-Microglia) were established by Draeger et al., 2022^27^. Briefly, iPSCs with stably integrated doxycycline-inducible transcription factors (PU.1, CEBPβ, IRF5, MAFB, CEBPα, and IRF8) and Trimethoprim (TMP)-inducible CRISPRi machinery – hereafter referred to as iTF-iPSCs – were maintained according to the above procedure. At day 0, iTF-iPSCs were replated onto Matrigel and PDL coated plates in Essential 8 Medium (Gibco; Cat. No. A1517001) supplemented with 10 µM ROCK inhibitor, 2 µg/mL doxycycline (Clontech; Cat. No. 631311), and 50 nM TMP (MP Biomedical; Cat. No. 195527). On day 2, the media was changed to Advanced DMEM/F12 Medium (Gibco; Cat. No. 12634-010) supplemented with 1L×LAntibiotic-Antimycotic (Anti-Anti) (Gibco; Cat. No. 15240-062), 1L×LGlutaMAX (Gibco; Cat. No. 35050-061), 2 μg/mL doxycycline, 50 nM TMP, 100 ng/mL Human IL-34 (Peprotech; Cat. No. 200-34) and 10 ng/mL Human GM-CSF (Peprotech; Cat. No. 300-03). On day 4, the media was changed to Advanced DMEM/F12 supplemented with 1L×LAnti-Anti, 1L×LGlutaMAX, 2 μg/mL doxycycline, 50 nM TMP, 100 ng/mL Human IL-34, 10 ng/mL, Human GM-CSF, 50 ng/mL Human M-CSF (Peprotech; Cat. No. 300-25), and 50 ng/mL Human TGFB1 (Peprotech; Cat. No. 100-21C). On day 8, the differentiated iTF-Microglia were used for assay or coculture.

### Human iPSC-derived neuron (iNeuron) differentiation

iPSC-derived neurons (iNeurons) were established by Tian et al., 2019^28^. Briefly, iPSCs with stably integrated doxycycline-inducible NGN2 – hereafter referred to as NGN2-iPSC s – were maintained according to the above procedure. On day -3, NGN2-iPSCs were replated onto Matrigel coated plates in Knockout DMEM/F-12 (Gibco; Cat. No. 12660012) supplemented with 1 x MEM Non-Essential Amino Acids Solution (Gibco; Cat. No. 11140-050), 1 x N2 supplement (Gibco; Cat. No. 17502-048), 10 ng/mL NT-3 (Peprotech; Cat. No. 450-03), 10 ng/mL BDNF (Peprotech; Cat. No. 450-02), 1 µg/mL mouse laminin (Gibco; Cat. No. 23017015), 10 µM ROCK inhibitor, and 2 µg/mL docycycline. On day -2, the cells were taken out of ROCK inhibitor by replacing the media with day –3 media formulation minus ROCK inhibitor. On day 0, the cells were washed with DPBS, lifted with Accutase, pelleted by 5 minutes of 300g centrifugation, and plated on PDL coated plates in BrainPhys Neuronal Medium (StemCell Technologies; Cat. No. 05790) supplemented with 0.5X N2 supplement, 0.5X B27 supplement (Gibco; Cat. No. 17504044), 10 ng/mL NT-3, 10 ng/mL BDNF, 1 µg/mL mouse laminin, and 2 µg/mL doxycycline. On day 3, the media was fully changed with day 0 media minus doxycycline. On day 7, half the media was removed and replaced by day 3 media.

### iNeuron and iTF-microglia coculture

iNeurons and iTF-microglia were differentiated as described above. Day 8 iTF-microglia were washed with DPBS, lifted with TrypLE Express (Gibco; Cat. No. 12605-028) for 10 minutes at 37 °C, quenched with Advanced DMEM/F-12, and pelleted by 8 minutes of 300g centrifugation. The supernatant was aspirated and the pellet was resuspended in day 0 neuronal media supplemented with 2X the cytokines and doxycycline used in day 4 microglia media. The iTF-microglia suspension was added to day 14 neurons to achieve a 10:3 (iNeuron:iTF-microglia) ratio in a half media change for a final concentration of 1X cytokines. Subsequent experiments occurred after three days of coculture.

### Lentivirus preparation

HEK293T cells were plated in tissue culture plastic dishes in DMEM (Gibco; Cat. No. 10313-039) supplemented with 10% fetal bovine serum (Gibco; Cat. No. A3382001) and 1% Penicillin-Streptomycin (Gibco; Cat. No. 15140122). Once 80-95% confluent, the media was fully changed in preparation for transfection. The transfection mix was prepared by mixing equal parts transfer DNA and Packaging Mix (0.8 µg psPAX2, 0.3ug pMD2G, 0.1 µg pAdVantage). This mixture was diluted with Opti-MEM I Reduced Serum Medium (Gibco; Cat. No. 31985070) before adding Lipofectamine 2000 Transfection Reagent (Invitrogen; Cat. No. 11668027). The solution incubated at room temperature for 10 minutes before being added to HEK293s. After two days, the media was taken up using a syringe and passed through a 0.45 µm filter. Lentivirus Precipitation Solution (Alstem; Cat. No. VC100) was added at a 1:4 (lentivirus precipitation solution:media) ratio, thoroughly mixed, and incubated at 4 °C overnight. The virus was pelleted by 30 minutes of 1500g centrifugation at 4 °C. The supernatant was removed and the pellet resuspended in DPBS.

#### Plasmid engineering

#### pKL078: pHIV CAG:Lck-mNeonGreen-P2A-NLS-mTagBFP2

pKL078 was prepared by ligating a gblock containing an Lck sequence and PCR amplified mNeonGreen and mTagBFP2 (Addgene plasmid #55320, mTagBFP2-PCNA-19-SV40NLS-4 was a gift from Michael Davidson^29^) products into a lentivirus backbone using Gibson assembly as per the manufacturer’s instructions.

#### pOT056: pCagg-Gamillus

pOT056 was prepared by PCR amplifying Gamillus (Gamillus/ pcDNA3 was a gift from Takeharu Nagai, Addgene plasmid #124837^30^) and ligating it into a lentivirus backbone using Gibson assembly as per the manufacturer’s instructions.

#### pOT020: pCagg-synaptophysin-Gamillus

pOT020 was prepared by PCR amplifying Gamillus (Gamillus/ pcDNA3 was a gift from Takeharu Nagai, Addgene plasmid #124837^30^) and synaptophysin (SynTagMA_pre was a gift from Thomas Oertner, Addgene plasmid #119738^31^) and ligating them into a lentivirus backboneusing Gibson assembly as per the manufacturer’s instructions.

### Membrane labeling of microglia

Lentivirus was prepared according to the above procedure for plasmid pKL078. iTF-iPSCs were kept according to the above procedure. Once confluent, the iTF-iPSCs were washed with DPBS, lifted using Accutase for 3 minutes at 37 °C, diluted with DPBS, and pelleted by 5 minutes of 220 g centrifugation. The supernatant was aspirated and the pellet was resuspended in StemFlex, 10 µM ROCK inhibitor, and lentivirus. The media was fully changed after two days. The transduced iTF-iPSCs were expanded, sorted on green fluorescence using the BD FACSAria Fusion (BD Biosciences), and maintained to be used for live imaging of iTF-microglia. To compare the morphology of monoculture and coculture iTF-microglia, these labeled microglia were plated 10:3 with unlabeled:labeled iTF-microglia or iNeurons:labeled iTF-microglia.

### Engineering neurons expressing Gamillus constructs

Lentivirus was prepared according to the above procedure for pOT056 and pOT020. NGN2-iPSCs were kept according to the above procedure. Once confluent, the NGN2-iPSCs were washed with DPBS, lifted using Accutase for 3 minutes at 37 °C, diluted with DPBS, and pelleted by 5 minutes of 220 g centrifugation. The supernatant was aspirated and the pellet was resuspended in StemFlex, 10 µM ROCK inhibitor, and lentivirus. The media was fully changed after two days. The transduced NGN2-iPSCs were expanded, sorted on green fluorescence using the BD FACSAria Fusion (BD Biosciences), and maintained to be used for live synaptic uptake assays.

### Quant-seq for characterization of iTF-microglia and iNeurons in coculture

Cocultures were prepared as previously described with membrane-labeled iTF-Microglia. Cell culture media was aspirated, cells washed twice with Hank’s Balanced Salt Solution (HBSS) (Gibco; Cat. No. 14025092), and lifted using Papain (Worthington Biochemical; Cat. No. LK003178) with 5 mM magnesium chloride and 5 µg/mL DNase (Worthington; Code: DPRF; Cat. No. LS006333). After 10 minutes at 37 °C, the papain was diluted with HBSS supplemented with 5 µg/mL DNAse and 5 mM magnesium chloride. The lifted cells were pelleted at 300g for 8 minutes. The supernatant was aspirated and the pellet resuspended in HBSS supplemented with 5 µg/mL DNAse and 5 mM magnesium chloride and immediately moved to ice. The cocultures were sorted into separate microglia and neuron populations by FACS based on the mNeonGreen expressed by the microglia.

Monocultured neurons were prepared as previously described. To compare to the coculture neurons, the monoculture neurons received a media change at day 14 containing IL-34, GM-CSF, TGFB, M-CSF, and dox according to the coculture media. Similarly, monocultured microglia were prepared such that at day 8, the microglia were lifted with TryplE and replated in the coculture media on PDL plates.

RNA from biological duplicates for each condition (more than 30,000 cells for each sample) was extracted using the Quick-RNA Miniprep Kit (Zymo; Cat. No. R1055). Libraries were prepared from total RNA (15Lng per sample) using the QuantSeq 3′ mRNA-Seq Library Prep Kit for Illumina (FWD) (Lexogen; Cat. No. 015UG009V0252) following the manufacturer’s instructions. Library amplification was performed with 17 total PCR cycles. mRNA-Seq library concentrations (mean of 5.36L±L2.37LngLμl^−1^) were measured with the Qubit dsDNA HS Assay Kit (Invitrogen; Cat. No. Q32851) on a Qubit 2.0 Fluorometer. Library fragment-length distributions (mean of 269L±L18Lbase pairs) were quantified with High Sensitivity D5000 Reagents (Agilent Technologies; Cat. No. 5067-5593) on the 4200 TapeStation System. The libraries were sequenced on an Illumina NextSeq 2000 P2 instrument with single-end reads.

Transcripts were aligned to the human reference genome (GRCh38, release 42) using Salmon v.1.4.0^32^ (the --noLengthCorrection, --validateMappings, --gcBias flags were used). Transcripts were mapped to genes and quantified using tximeta. To visualize gene expression for genes related to neurons, microglia, complement, and microglial activation, the row centered and scaled counts were displayed using Enhanced Heaptmap (gplots, version 3.1.3.1).

### Flow cytometry assay for uptake of Gamillus protein

NGN2-iPSCs engineered with various Gamillus constructs were differentiated according to the above protocol. iTF-iPSCs were transduced with a non-targeting sgRNA that includes a nuclear BFP marker, differentiated, and cocultured with the aforementioned engineered neurons. After three days of coculture, the cells were washed with HBSS, lifted using Papain with 5 mM magnesium chloride and 5 µg/mL DNase for 10 minutes at 37 °C, and diluted with HBSS. Single cell fluorescence was measured by flow cytometry using the BD FACS Celesta (BD Biosciences). Microglia uptake of Gamillus was analyzed by measuring the median green fluorescence of the BFP+ population. For synaptic uptake experiments with pharmacological treatments, cocultures were treated with 200 nM Cytochalasin D (Invitrogen; Cat. No. PHZ1063). For annexin V treatments, day 14 neurons were incubated with 1:100 annexin V (Invitrogen; A13202) for 15 minutes at 37 °C, washed with HBSS, and replenished with neuron conditioned media before iTF-Microglia were added for coculture.

### Library preparation and synaptic pruning screen

Top and bottom oligos for two sgRNAs targeting each of the 102 ASD-risk genes^16^ and 28 non-targeting controls were ordered from IDT (Supplementary Table 1). Sequences were chosen according the two highest predicted activity scores from Horlbeck et al., 2016^33^. Top and bottom oligos were annealed then pooled before ligating into the sgRNA backbone, pMK1334^28^, that contains a nuclear BFP. Lentivirus was prepared according to the previously described protocol. iTF-iPSCs were transduced with the library with MOI < 0.15, selected using 1 µg/mL Puromycin dihydrochloride (Sigma; Cat. No. P9620), and differentiated into microglia for monoculture phagocytosis of beads (Polysciences; Cat. No. 15702-10) and synaptosomes or for coculture with iNeurons expressing synaptophysin-Gamillus. Synaptosomes were prepared using iNeurons expressing the synaptophysin-Gamillus construct and SynPer (Thermo Scientific; Cat. No. 87793) following the manufacturer’s protocol. Microglia were treated with 3 mg/mL of synaptosomes or 1:1,000 dilution of beads. After 2 hours of incubation with beads or synaptosomes and 3 days of coculture with iNeurons expressing the synaptophysin-Gamillus, cells were washed with HBSS then lifted with Papain with 5 mM magnesium chloride and 5 µg/mL DNase for 10 minutes at 37 °C. Cells were diluted with HBSS and pelleted before resuspension in HBSS with 5 mM magnesium chloride and 5 µg/mL DNase. Microglia (BFP+ cells) were sorted by FACS (BD FACSAria Fusion) into populations with low (bottom 25%) or high (top 25%) Gamillus-FITC signal. Low and high Gamillus-FITC samples from three replicates were prepared for next-generation sequencing as previously described^27^. Briefly, genomic DNA was isolated using a Macherey-Nagel Blood L kit (Macherey-Nagel; Cat. No. 740954.20) and sgRNA sequences were amplified and sequenced on an Illumina MiSeq V3.

Subsequent sgRNA quantifications were completed using a new bioinformatics pipeline^34^ (https://github.com/noamteyssier/sgcount). Briefly, sequencing reads were mapped to protospacer sequences in our library (Supplementary Table 1). sgRNA counts were then analyzed using an updated MAGeCK-iNC pipeline^28^ that uses a negative binomial distribution (https://noamteyssier.github.io/crispr_screen). Phenotypes and p-values for each gene were calculated using a new algorithm, geopagg (https://github.com/noamteyssier/geopagg), that uses Benjamini-Hochberg correction for multiple hypothesis correction, an empirical false discovery rate (FDR), and a weighted geometric mean of individual guides contributing to an aggregated gene FDR and fold change. Hit genes were called based on an FDR of 0.01.

### qPCR

To validate knockdowns for *TREM2*, *ADNP, TCF4,* and *TCF7L2*, RNA was extracted using the Quick-RNA Microprep Kit (Zymo Research, Cat. No. R1050) with DNA digest. RNA was converted to cDNA using SensiFAST cDNA Synthesis Kit (Meridian Bioscience, Cat. No. BIO-65053). qPCR reactions were setup using SensiFAST SYBR Lo-ROX 2X Master Mix (Bioline; Cat. No. BIO-94005), custom qPCR primers from Integrated DNA Technologies used at a final concentration of 0.2µM, and 5 ng of cDNA. Reaction amplification was measured using the Applied Biosystems QuantStudio 6 Pro Real-Time PCR System using QuantStudio Real Time PCR software (v.1.3). Expression fold changes were calculated using the ΔΔCt method, normalizing to housekeeping gene *GAPDH*.

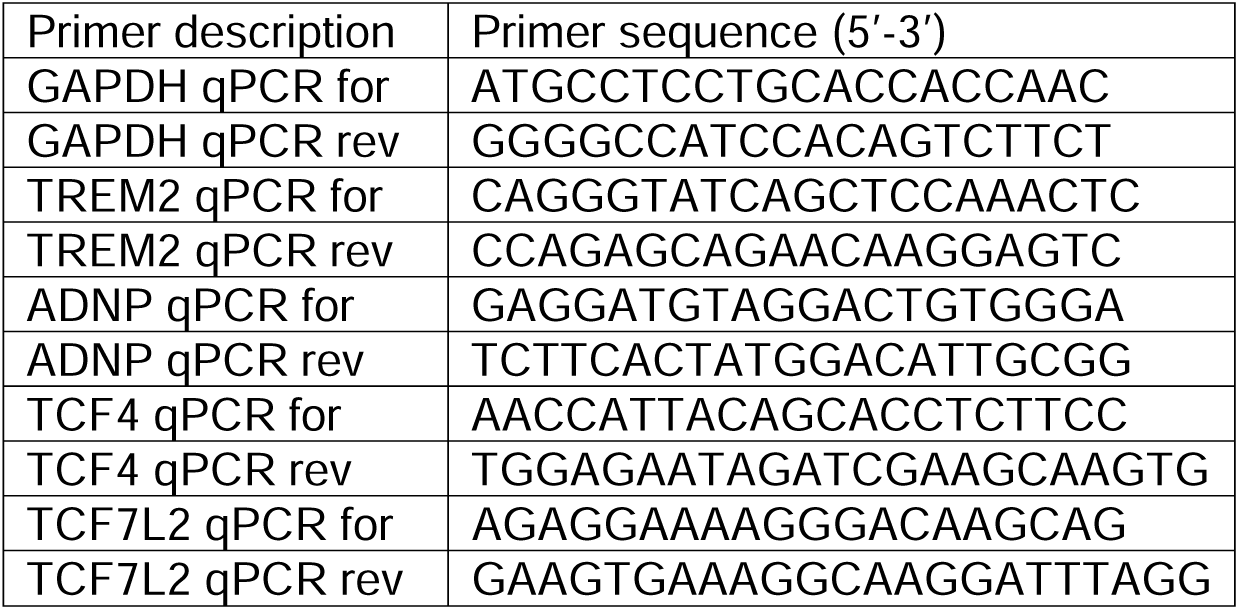

### Immunohistochemistry (IHC)

Cell cultures were first fixed using 4% paraformaldehyde (Electron Microscopy Sciences; Cat. No. 15710) for 15 minutes at room temperature. Cells were then washed thrice with DPBS before blocking and permeabilizing for one hour at room temperature with blocking buffer: DPBS supplemented with 5% Normal Goat Serum (Vector Laboratories; Cat. No. S-1000-20) with 0.01% Triton X-100 (TEKnova; Cat. No. T1105). Primary antibodies were in blocking buffer and left at 4 °C overnight. Samples were washed thrice with DPBS. Secondary antibodies were diluted in blocking buffer and added for 1-2 hours at room temperature. Secondary antibody solutions were washed thrice with DPBS before imaging using the Molecular Devices Image Express Confocal HT.ai or IN Cell Analyzer 6000 (GE; Cat. No. 28-9938-51).

| Target | Dilution | Catalog Number |
| --- | --- | --- |
| Anti-ADNP | 1:500 (IF), 1:1000 (WB) | Novus Biologicals; NBP1-89236 |
| Anti-CLTC | 1:500 (IF) | ProteinTech; 66487-1-Ig |
| Anti-Rab5A | 1:200 (IF) | Cell signaling; 46449 |
| Anti-Rab11A | 1:200 (IF) | Invitrogen; MA5-49197 |
| Anti-CD68 | 1:1000 (IF) | BioRad; MCA5709 |
| Anti-GM130 | 1:500 (IF) | Abcam; ab169276 |
| Anti-COL1A1 | 1:200 (IF) | Cell Signaling; 66948 |
| Anti-α Tubulin | 1:1000 (IF) | Cell Signaling; 3873 |
| Anti-NeuN | 1:500 (IF) | Abcam; ab279296 |
| Anti-GAPDH | 1:50,000 (WB) | SCBT; sc-47724 |
| Anti-vGlut1 | 1:500 (IF) | Sigma; MAB5502 |
| Anti-Homer | 1:500 (IF) | Synaptic Systems; 160003 |

### Surveillance dynamics

Membrane labeled iTF-Microglia were cocultured with iNeurons. After three days of coculture, time lapse images were acquired using a 20X objective every 15 seconds for 30 minutes on an IN Cell Analyzer 6000 (GE; Cat. No. 28-9938-51). Images were then segmented on the membrane label using CellProfiler’s IdentifyPrimaryObjects. Segmentations for a given field of view were stacked and stabilized using ImageJ. Individual cells from various fields were cropped for single cell analysis of total area surveilled which equated to total area of the segmentation stack’s max projection normalized to initial cell area.

### Phagocytosis assays with in-well controls

The sgRNA backbone, pMK1334^28^, that contains a nuclear BFP was modified using Gibson assembly to create a backbone containing a nuclear mApple protein. A non-targeting control (NTC 2) protospacer sequence was ligated into the new nuclear mApple backbone and protospacer sequences targeting *TREM2*, *ADNP, TCF4,* and *TCF7L2* were ligated into the nuclear BFP backbone. Uptake of synaptic material with the in-well NTC iTF-Microglia was conducted as previously described with the modification that half of the microglia in the well carried the NTC sgRNA with nuclear mApple and the other half a gene targeting sgRNA with nuclear BFP. The same mixture of microglia was plated in monoculture for phagocytosis of synaptosomes and fluorescent beads (Polysciences; Cat. No. 15702-10). Synaptosomes were prepared using iNeurons expressing the synaptophysin-Gamillus construct and SynPer (Thermo Scientific; Cat. No. 87793) following the manufacturer’s protocol. Microglia were treated with 3 mg/mL of synaptosomes or 1:1,000 dilution of beads for 1 hour then lifted using TryplE for 10 minutes at 37 °C. Single-cell fluorescence was measured by flow cytometry using the BD FACS Celesta (BD Biosciences).

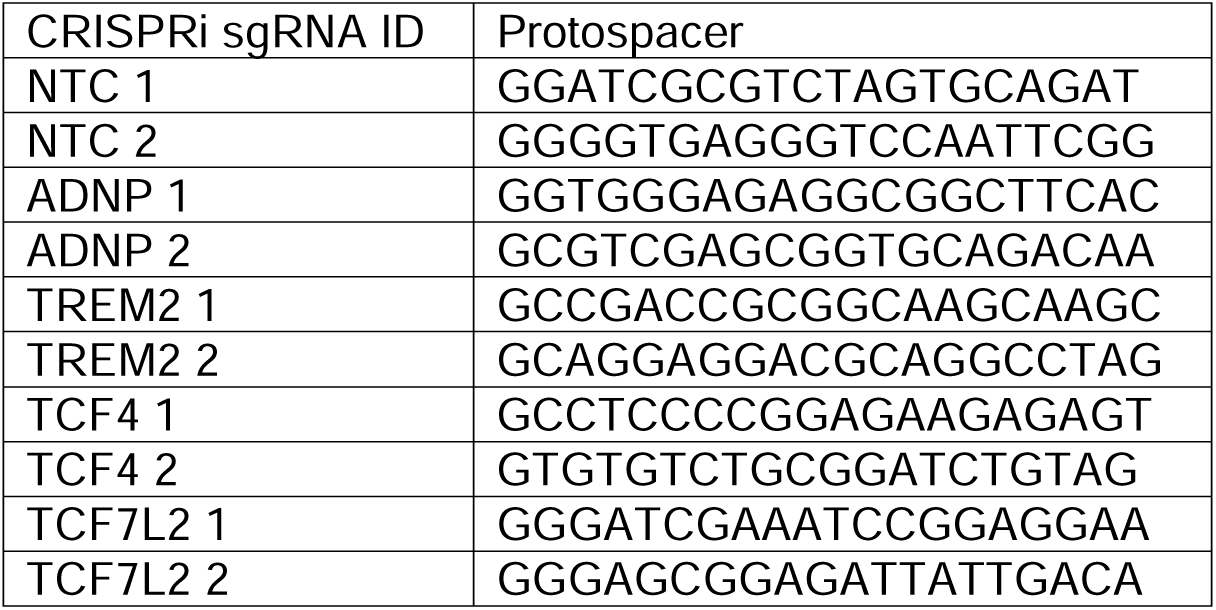

### CROPseq analysis

Single cell RNA-sequencing of iTF-Microglia with CRISPRi-based gene knockdowns of 53 ASD risk genes was conducted and analyzed according to the methods described in Dräger 2022^27^. In brief, Cell Ranger (v.5.0.1, 10X Genomics) was used for alignment to the GRCh38-3.0.0 reference genome and gene expression quantification. DemuxEM71 was used for sgRNA assignment. The resulting data represented 4,466 cells and an average of 17,120 reads per cell (Supplementary Figure 3A-D). Differential expression analysis comparing single cells with sgRNAs targeting *ADNP* to NTCs was performed using the resulting Seurat object and FindMarkers function.

### Surface proteomics

Surface proteomics was performed according to the protocol established in Kirkemo et al., 2022^35^. In brief, cells were lifted using Versene for 10 minutes at 37 °C. Cells were diluted in DPBS and pelleted by centrifugation at 1500 g for 5 minutes and resuspended in DPBS pH 6.5. Cell surfaces were labeled by incubating 0.5 µM of WGA-HRP for 5 minutes on ice, then adding 500 µM biotin tyramide, and finally adding 1 mM of H2O2. The mixture was incubated at 37 °C for 2 minutes. The reaction was quenched with 10 mM Sodium Ascorbate/1 mM Sodium Pyruvate followed by two additional washes in the same buffer prior to a final wash in PBS. Cells were pelleted and flash frozen before mass spectrometry.

The cell pellet was thawed and processed for LC-MS/MS using a preOmics iST kit (P.O. 00027). Briefly, cell pellets were lysed in 2x RIPA buffer containing protease inhibitors (cOmplete, Mini, EDTA-free Protease Inhibitor Cocktail; Millipore Sigma 11836170001) and 1.25 mM EDTA. Cells were lysed via sonication. Biotinylated proteins were pulled down with NeutrAvidin-coated agarose beads (Thermo) for 1 hour at 4 °C. Beads were transferred to Poly-Prep chromatography columns (Bio-Rad) and sequentially washed with 1x RIPA buffer, high-salt PBS (PBS pH 7.4, 2 M NaCl [Sigma]), and denaturing urea buffer (50 mM ammonium bicarbonate, 2 M Urea). From the PreOmics iST kit, 50 µL of the provided LYSE solution was added to the slurry and the mixture was incubated at 55 °C for 10 minutes with shaking. The provided enzymes mixture (Trypsin and LysC) were resuspended in 210 µl of RESUSPEND buffer, mixed, and added to the slurry. Samples were allowed to mix at 500 rpm for 1.5 hours at 37 °C, before being quenched with 100 µl of STOP solution. Sample was spun in provided C18 spin cartridge and washed 1X with 200µl of WASH 1 and WASH 2. Peptides were eluted with 2X 100 µl of ELUTE, dried, and resuspended with 2% acetonitrile and 0.1% TFA. Peptides were quantified using Pierce Quantitative Colorimetric Peptide Assay (Thermo Fisher Scientific, 23275).

Mass spectrometry was performed as described previously in Kirkemo et al., 2022^35^. Briefly, 200 ng of samples were separated over a 90 minute 3%-35% ACN gradient on either an Aurora Ultimate CSI 25cm×75µm C18 (Ionopticks) or a PepSep XTREME 25 cm x 150 µm (Bruker) UHPLC column using a nanoElute UHPLC system (Bruker), and injected into a timsTOF pro mass spectrometer (Bruker). For database searching, peptides were searched using PEAKS Online X version 1.7 against both the plasma membrane annotated human proteome (Swiss-prot GOCC database, August 3, 2017 release). Enzyme specificity was set to trypsin + LysC with up to two missed cleavages. Cysteine carbamidomethylation was set as the only fixed modification; acetylation (N-term) and methionine oxidation were set as variable modifications. The precursor mass error tolerance was set to 20 PPM and the fragment mass error tolerance was set to 0.05 Da. Data was filtered at 1% for both protein and peptide FDR and triaged by removing proteins with fewer than 2 unique peptides. All mass spectrometry database searching was based off three biological replicates. Biological replicates underwent washing, labeling, and downstream LC-MS/MS preparation separately.

### Viability assay

Live cell cultures were incubated with TO-PRO-3 (Invitrogen; Cat. No. T3605) and Hoechst 33342 (Thermo Fisher Scientific; Cat. No. H3570) at 37 °C for 15 minutes. Cultures were then imaged and nuclei were classified using CellProfiler pipelines as live or dead with dead cells having a high average TO-PRO-3 intensity.

### Western blotting

Cells were washed twice with cold DPBS then lysed using 1 X RIPA buffer (Abcam; Cat. No. 156034) with protease (Roche; Cat. No. 11836170001) and phosphatase inhibitors (Roche; Cat. No. 4906845001). Cells lysates were collected and centrifuged at max speed (16,000 xG) for 5 minutes at 4 °C. Protein concentration of the supernatant was measured using a BCA Protein Assay Kit (Pierce; Cat. No. 23225). 10 µg of protein was mixed with 1 X Laemmli SDS-Sample buffer, Reducing (bioworld #10570021-2) and boiled for 5 minutes at 95 °C. Denatured proteins were loaded into a 4-12% Bis-Tris polyacrylamide gel (Invitrogen; Cat. No. NP0336BOX) and run at 120 V for 1 hour. Proteins were transferred to a nitrocellulose membrane using BioRad Transblot Turbo Transfer System for 11 minutes at 25 V. Nitrocellulose membranes were blocked for 1 hour at room temperature with Intercept (PBS) blocking buffer (Licor; Cat. No. 927-70001). Primary antibodies diluted in Intercept (PBS) blocking buffer were added overnight at 4 °C. Membranes were washed thrice with TBST. Secondary antibodies diluted in Intercept (PBS) blocking buffer were added for 2 hours at room temperature. Membranes were washed thrice with TBST before being imaged on a Licor Immunoblots.

### Endolysosomal characterization

Cells were pretreated with 30 μM Dynasore (Sigma; Cat. No. 324410) or 200 nM Bafilomycin A (Sigma; Cat. No. SML1661) for 2 hours. Dextran, Alexa Fluor488 10,000 MW (Invitrogen; Cat. No. D22910) was diluted to 25 μg/mL in day 4 microglia media, added to cell cultures, and incubated at 37 °C for 30 minutes. pHrodo Red dextran10,000 MW (Invitrogen; Cat. No. P10361) was diluted to 10 μg/mL in day 4 microglia media, added to cell cultures, and incubated at 37 °C for 2 hours. Lysotracker Deep Red (Invitrogen; Cat. No. L12492) was diluted to 20 nM in day 4 microglia media, added to cell cultures, and incubated at 37 °C for 30 minutes. Lysosensor Green (Invitrogen; Cat. No. L7535) was diluted to 500 nM in day 4 microglia media, added to cell cultures, and incubated at 37 °C for 30 minutes. DQ Green BSA (Invitrogen; Cat. No. D12050) was diluted to 10 μg/mL in day 4 microglia media, added to cell cultures, and incubated at 37 °C for various times. After incubation, iTF-Microglia were first imaged using an IN Cell Analyzer 6000 (GE; Cat. No. 28-9938-51) then washed and lifted using TryplE at 37 °C for 10 minutes. Cells were diluted in cold HBSS before measurement by flow cytometry using the BD FACS Celesta (BD Biosciences).

### Cytokine immunoassay

Secreted proteins in iTF-Microglia conditioned media were measured using the Proteome Profiler Human Cytokine Array Kit (R&D Systems; Cat. No. ARY005B) following the manufacturer’s protocol. Chemiluminescent images were acquired using ChemiDoc Touch Imaging System. Integrated pixel intensity for each dot was measured using ImageJ. Integrated pixel intensities for technical replicate dots were averaged before subtracting the background as measured by the integrated pixel intensity for negative dots.

### TREM2 Homogeneous Time Resolved Fluorescence (HTRF) assay

TREM2 concentrations were measured using the HTRF Human TREM-2 Detection Kit (Revvity; Cat. No. 63ADK099PEG) following the manufacturer’s instructions. Cell lysates were collected using 1 X RIPA buffer with protease and phosphatase inhibitors. Cell lysate and media TREM2 concentrations were determined according to standard curves prepared in the aforementioned RIPA buffer or unconditioned media, respectively.

## RESULTS

### A subset of ASD risk genes are expressed by microglia and their perturbation shifts the activation state of iPSC-derived microglia (iTF-Microglia)

Although the expression of ASD risk genes is enriched in excitatory neurons^16^, we asked whether a subset of ASD risk genes may be meaningfully expressed by microglia. Using a published human single-cell sequencing dataset^36^, we plotted the expression of 102 ASD risk genes^16^ by neurons compared to the expression by microglia (Figure 1A). Some genes showed much higher, nearly exclusive expression by neurons, but many genes showed comparable levels of expression by both cells and some genes highly expressed by microglia. Those genes that are highly expressed by neurons relate to synaptic function while those that are comparably or highly expressed by microglia relate to gene regulatory pathways. To understand the impact of regulatory genes in microglia, we used single-cell sequencing combined with CRISPRi-based gene knockdowns (CRISPR droplet sequencing, CROP-Seq) in human iPSC-derived microglia (iTF-Microglia) (Figure 1B). Our sgRNA library targeted 53 genes, using two guides per gene, that are highly expressed in our iTF-Microglia. We found that over 95% of sequenced cells expressed the microglia identity marker, Iba1, encoded by *AIF1* (Figure 1C). Cluster analysis revealed five transcriptomic states occupied by the iTF-Microglia: chemokine state (enriched for *CCL13*), interferon state (enriched for *IFIT1*), proliferative state (enriched for *MKI67*), and two non-microglial states that largely do not express Iba1 and are driven by *KMT2C* and *TAOK1* knockdown (Figure 1D, Supplemental Figure 1D,E). We analyzed the UMAP distribution (Figure 1E,F) and cluster occupancy (Figure 1G-I) of the 12 most represented ASD risk gene knockdowns in the CROP-Seq dataset (Supplemental Figure 1A-C). Many ASD risk gene knockdowns – notably *ADNP*, *AP2S1*, *TRIP12,* and *ZMYND8* – altered the composition of microglia activation states. Our transcriptomic characterization suggests altered activation states for iTF-Microglia induced by some ASD risk gene knockdowns. We also observed an anti-correlation between changes in the interferon and chemokine states such that gene knockdowns that increase occupancy in the interferon state decrease occupancy in the chemokine state (Figure 1I, Supplemental Figure 1F). We next asked whether ASD risk gene knockdowns in iTF-Microglia and their corresponding transcriptional changes cause functional impacts.

**Figure 1.**
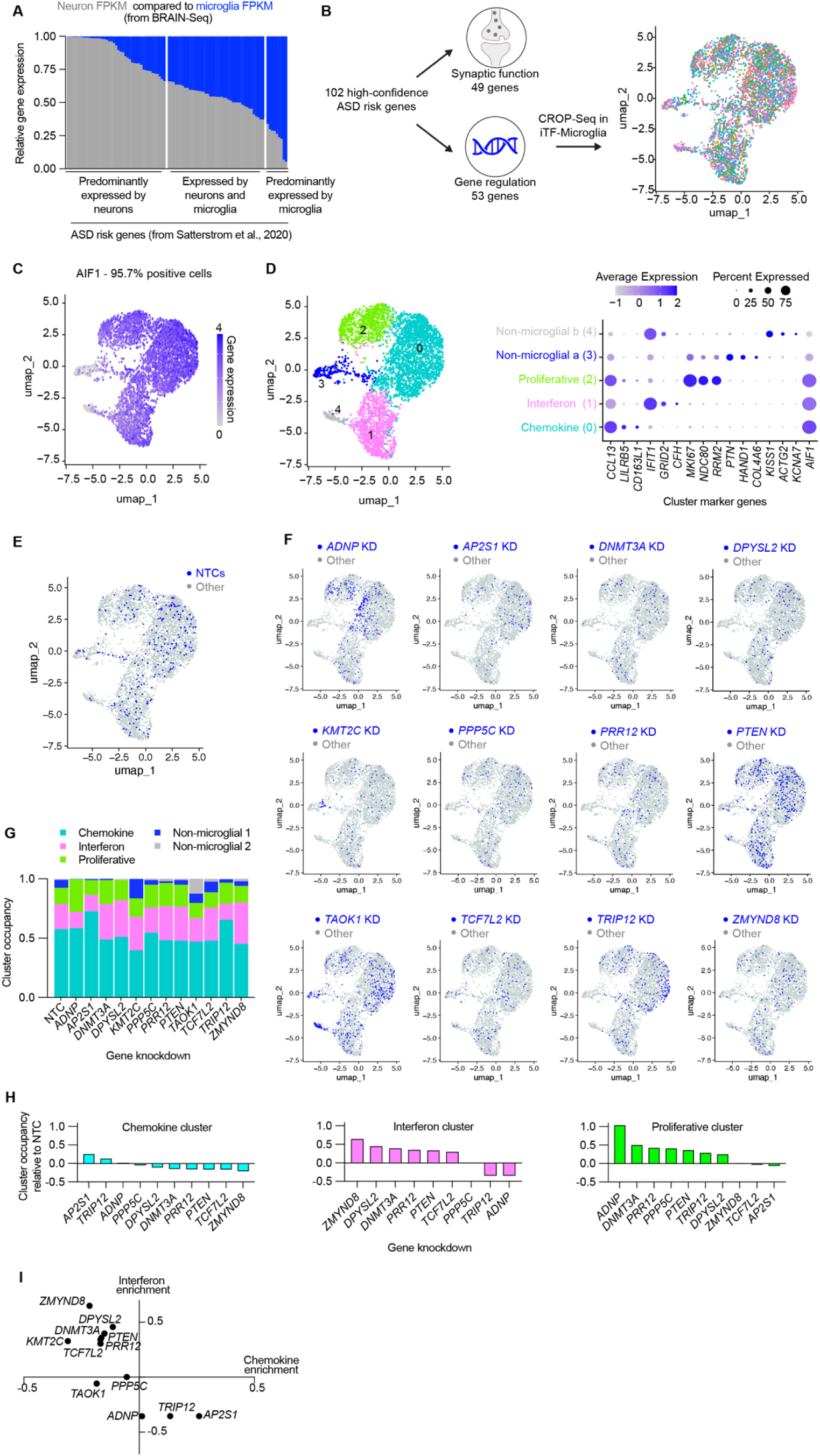
A subset of ASD risk genes are expressed by microglia and their perturbation shifts the activation state of iPSC-derived microglia (iTF-Microglia). A. Stacked bar graph comparing the relative expression of 102 ASD risk genes^16^ in neurons and microglia. B. UMAP derived from single cell transcriptomic data for iTF-Microglia with CRISPRi-mediated knockdowns targeting 53 ASD risk genes involved in gene regulation. Each dot represents a single cell colored by gene target. C. UMAP displaying the expression of the microglia marker, IBA1, encoded by *AIF1* for individual cells. D. (Left) UMAP displaying the results of transcriptomic cluster analysis for which each color represents a unique cluster. (Right) Dot plot showing the expression of each cluster’s top three differentially expressed genes. The color of the dots reflect the relative expression of the gene in a given cluster, and the size of the dots represent the percent of cells expressing the gene in a given cluster. E-F. UMAPs displaying the distribution of cells containing sgRNAs targeting 12 specific ASD risk genes or non-targeting controls (NTCs). Blue dots represent the target cell and gray dots represent all other cells. G. Stacked bar graph comparing the cluster occupancy by cells containing sgRNAs targeting 12 specific ASD risk genes or NTCs. Bar segments are colored according to cluster identity. H. Individual bar graphs for the three activation clusters showing the change in cluster occupancy relative to NTCs. I. Scatterplot showing the change in the interferon and chemokine clusters relative to NTCs.

### Developing a high-throughput, *in vitro* assay for synaptic pruning by iPSC-derived microglia (iTF-Microglia)

Microglial remodeling of synapses is an important function during neurodevelopment^11,13,37^. Given the observations of altered synapse density in ASD, it is speculated that dysregulated microglial synaptic pruning may contribute to ASD biology^3^. We set out to systematically assess the role of ASD risk genes in iTF-Microglial synaptic pruning. The standard method for measuring synaptic pruning involves volumetric confocal microscopy and advanced image analysis techniques to count synaptic puncta colocalized within microglial lysosomes. This method is not compatible with performing a high-throughput CRISPRi-based screen, so we developed a flow cytometry-based readout for synaptic pruning (Figure 2A).

**Figure 2.**
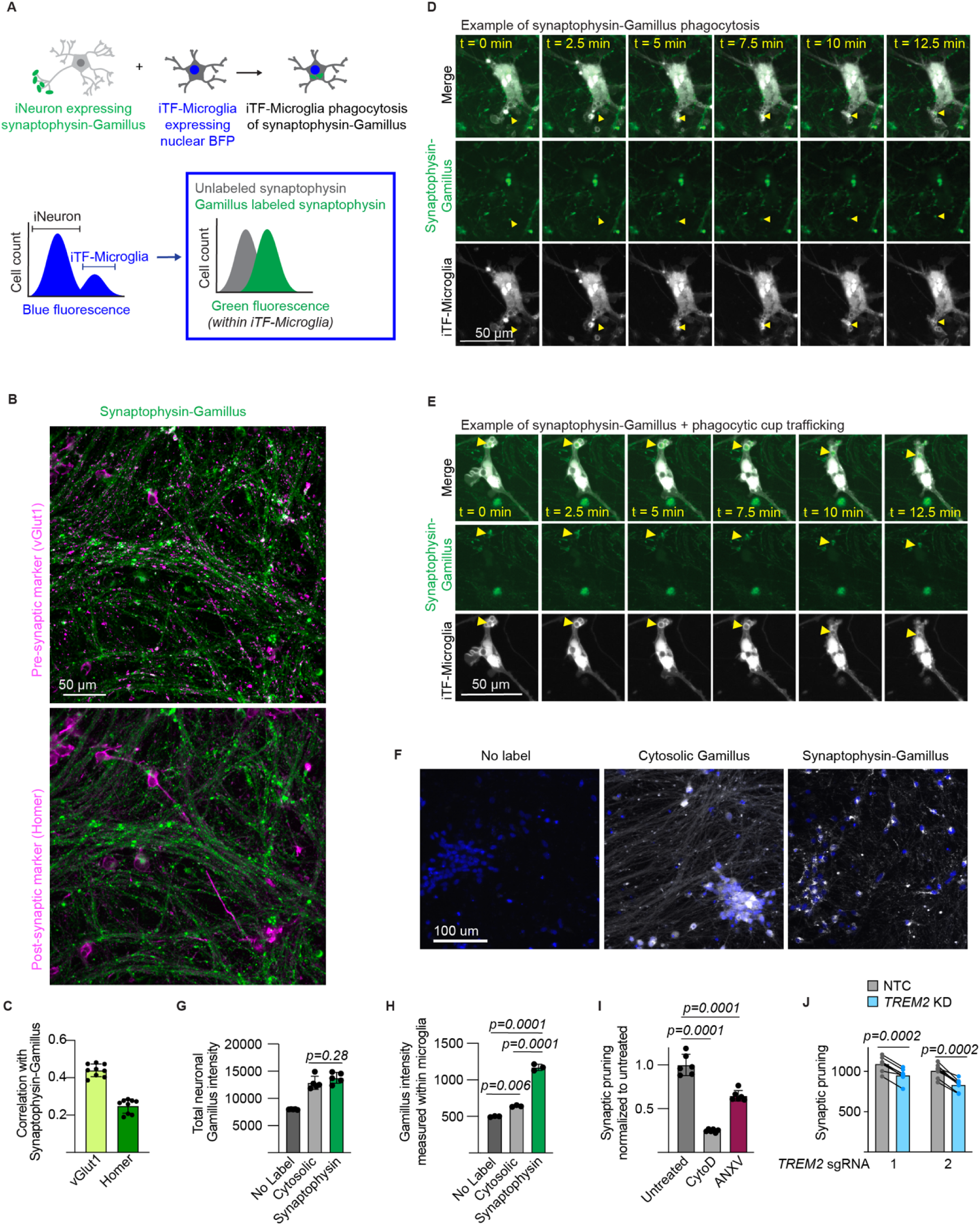
iPSC-derived microglia (iTF-Microglia) and iPSC-derived neuron (iNeuron) coculture enables monitoring of microglial synaptic pruning. A. Experimental and analytical strategy for measuring iTF-Microglia uptake of synaptic material from iPSC-derived neurons (iNeurons) using flow cytometry. (Top) iNeurons are engineered to express synaptophysin linked to the acid-tolerant green fluorescent protein, Gamillus. iTF-Microglia are engineered to express a nuclear blue fluorescent protein. (Bottom) After coculture and measurement using flow cytometry, iTF-Microglia are identified by their blue fluorescence, and the amount of green fluorescence from uptake of synaptophysin-Gamillus is measured within this iTF-Microglia population. B. Representative micrographs of iNeurons expressing synaptophysin-Gamillus (green) costained with an antibody against a presynaptic marker, vGlut1 (magenta, top), or a post-synaptic marker, Homer (magenta, bottom). C. Correlation analysis of synaptophysin-Gamillus with vGlut1 or Homer (N = 3 fields of view from 3 wells, bars represent mean +/- standard deviation). D,E. Time lapse images of iNeurons expressing synaptophysin-Gamillus (green) and iTF-Microglia expressing a membrane marker, Lck-mApple (gray). D. The yellow arrow highlights a synaptophysin-Gamillus punctum that is taken up into a phagocytic cup and beginning to be trafficked toward the iTF-Microglia’s soma. E. The yellow arrow highlights a synaptophysin-Gamillus puncta inside a phagocytic cup that is being trafficked toward the iTF-Microglia’s soma. F. Representative micrographs of iNeurons engineered to express cytosolic Gamillus or synaptophysin-Gamillus. Nuclei are marked by Hoechst 33342 and displayed in blue. Gamillus is displayed in gray. G. Expression of the Gamillus protein measured by total fluorescence intensity per field of view (n = 5 wells, bars represent mean +/- standard deviation, Tukey’s multiple comparisons test). H. Uptake of Gamillus by iTF-Microglia in coculture measured by flow cytometry (n = 3 wells, bars represent mean +/-standard deviation, Tukey’s multiple comparisons test). I. Uptake of synaptophysin-Gamillus by iTF-Microglia in cocultures untreated, treated with actin polymerization inhibitor, Cytochalasin D, or treated with phosphatidylserine blocker, annexin V, as measured by flow cytometry (n = 5 wells, bars represent mean +/- standard deviation, Tukey’s multiple comparisons test). J. Uptake of synaptophysin-Gamillus by iTF-Microglia with *TREM2* targeting sgRNAs compared to uptake of synaptophysin-Gamillus by in-well iTF-Microglia with non-targeting control (NTC) sgRNA (n = 5 wells, bars represent mean, connected dots represent NTC and *TREM2* knockdown microglia in the same well, multiple paired t-tests).

First, we established a scalable and rapid coculture protocol such that iTF-Microglia and iNeuron interactions can be assessed within 17 days (Supplementary Figure 2A). The cocultured cells retain their identity and iTF-Microglia show a trend towards decreased activation as measured by RNA-Sequencing (Supplementary Figure 2B). Towards developing a readout for synaptic pruning within the coculture, iNeurons were engineered with a presynaptic protein, synaptophysin, linked to an acid-tolerant, green fluorescent protein, Gamillus, and iTF-Microglia were engineered with a nuclear blue fluorescent protein. Upon coculture, the microglia population can be identified based on high blue fluorescence and the green fluorescence associated with uptake of synaptophysin-Gamillus from neurons can be measured (Figure 2A).

We first characterized our proposed strategy by investigating neuronal expression of the synaptophysin-Gamillus. iNeurons engineered with synaptophysin-Gamillus were stained for a presynaptic protein, vGlut1, and a post-synaptic protein, Homer (Figure 2B). Correlation analysis indicated that synaptophysin-Gamillus better correlated with vGlut1 compared to Homer, suggesting a preferentially pre-synaptic localization of synaptophysin-Gamillus (Figure 2C). A limitation of this construct is that a portion of synaptophysin-Gamillus was localized to the neuronal soma. However, time lapse imaging revealed that some somatic synaptophysin-Gamillus puncta were trafficked down neurites (Supplementary video 1). Time lapse imaging of neurons expressing synaptophysin-Gamillus cocultured with membrane-labeled iTF-Microglia revealed uptake and internal trafficking of synaptophysin-Gamillus puncta from neurites by iTF-Microglia (Figure 2D,E).

To assess the selectivity for uptake of specific types of neuronal materials by iTF-Microglia using this strategy, we compared the uptake of fluorescent material in iTF-microglia cocultured with neurons expressing synaptophysin-Gamillus to those cocultured with neurons expressing untagged Gamillus, which shows a diffuse cytosolic localization (Figure 2F-H, Supplementary Figure 2C). The expression of the Gamillus protein as measured by total fluorescence intensity within the iNeurons did not differ between the two engineered iNeuron cultures (Figure 2G). After coculturing the two engineered iNeuron populations with iTF-Microglia and quantifying the uptake of fluorescent material by the iTF-Microglia, we saw a significantly greater uptake of fluorescent material from neurons expressing synaptophysin-Gamillus compared to cytosolic Gamillus, suggesting that iTF-microglia selectively take up synaptic material in our assay (Figure 2H, Supplementary Figure2D-E). The uptake of cytosolic Gamillus may be attributed to the clearance of dead neurons by iTF-Microglia as we observe an increase in culture viability and decrease in cell counts upon coculture (Supplementary Figure 2F,G).

To further benchmark our proposed strategy, we used pharmacologic and genetic approaches. We treated cocultures with an actin polymerization inhibitor, Cytochalasin D. The large decrease in synaptophysin-Gamillus intensity within iTF-Microglia upon Cytochalasin D treatment suggests that the readout is relaying a phagocytosis-dependent process (Figure 2I, Supplementary Figure3A). We further confirmed that the measured fluorescence intensity is indeed from internalized synaptophysin-Gamillus by measuring uptake after increasing incubation times with lysosome acidification inhibitor, Bafilomycin A, and measuring uptake in the presence of a cell impermeable, green fluorescence quencher, Trypan Blue (Supplementary Figure 3B,C). Uptake of synaptic material has been shown to be dependent on externalized phosphatidylserine^38,39^. We treated cocultures with a phosphatidylserine blocker, annexin V. We also see a decrease in synaptophysin-Gamillus intensity within iTF-Microglia upon annexin V treatment, suggesting that the readout is also phosphatidylserine-dependent (Figure 2I, Supplementary Figure 3D). In developmental contexts, TREM2 has been shown to mediate synaptic pruning^37^. In our *in vitro* system, we see that knockdown of *TREM2* in iTF-Microglia reduces uptake of synaptophysin-Gamillus (Figure 2J, Supplementary Figure 3E). Altogether, we designed, engineered, and characterized a high-throughput, *in vitro* assay to measure iTF-Microglial synaptic pruning to be used in a CRISPRi-based screen of ASD-risk genes.

### Convergent and divergent phagocytosis phenotypes reveal *ADNP* as a unique genetic regulator of iPSC-derived microglia (iTF-Microglia)

We conducted three phagocytosis CRISPRi-based screens using beads and iNeuron-derived synaptosomes as counter screens to synaptic pruning to ultimately assess the functional impact of 102 ASD risk genes in iTF-Microglia (Figure 3A). We delivered sgRNAs targeting 102 ASD risk genes^16^ with two guides per gene and 28 non-targeting controls to iPSCs stably expressing the CRISPRi machinery^27^ that were then differentiated into iTF-Microglia to produce microglia with knockdowns (KDs) of individual ASD risk genes (Supplementary Figure 4A). Beads and iNeuron-derived synaptosomes were added to microglia cultures. For synaptic pruning, the microglia were cocultured with the iNeurons expressing synaptophysin-Gamillus. For all screens, we used fluorescence-activated cell sorting (FACS) to sort iTF-Microglia into two populations: one with high green fluorescence (top 25%) or high phagocytosis of the given substrate and one with low green fluorescence (bottom 25%) or low phagocytosis of the given substrate. By comparing the frequencies of cells expressing each sgRNA across the two populations as determined by next-generation sequencing and comparing them to non-targeting control sgRNAs (NTCs), we identified genes that significantly increased or decreased phagocytosis of beads, iNeuron-derived synaptosomes, and synaptic pruning when knocked down (Figure 3B-D). We did not find any correlation between the basal expression, extent of knockdown, or classification as hit or non-hit amongst target genes (Supplementary Figure 4B-D) suggesting that the screen results are not confounded by technical limitations of CRISPRi.

**Figure 3.**
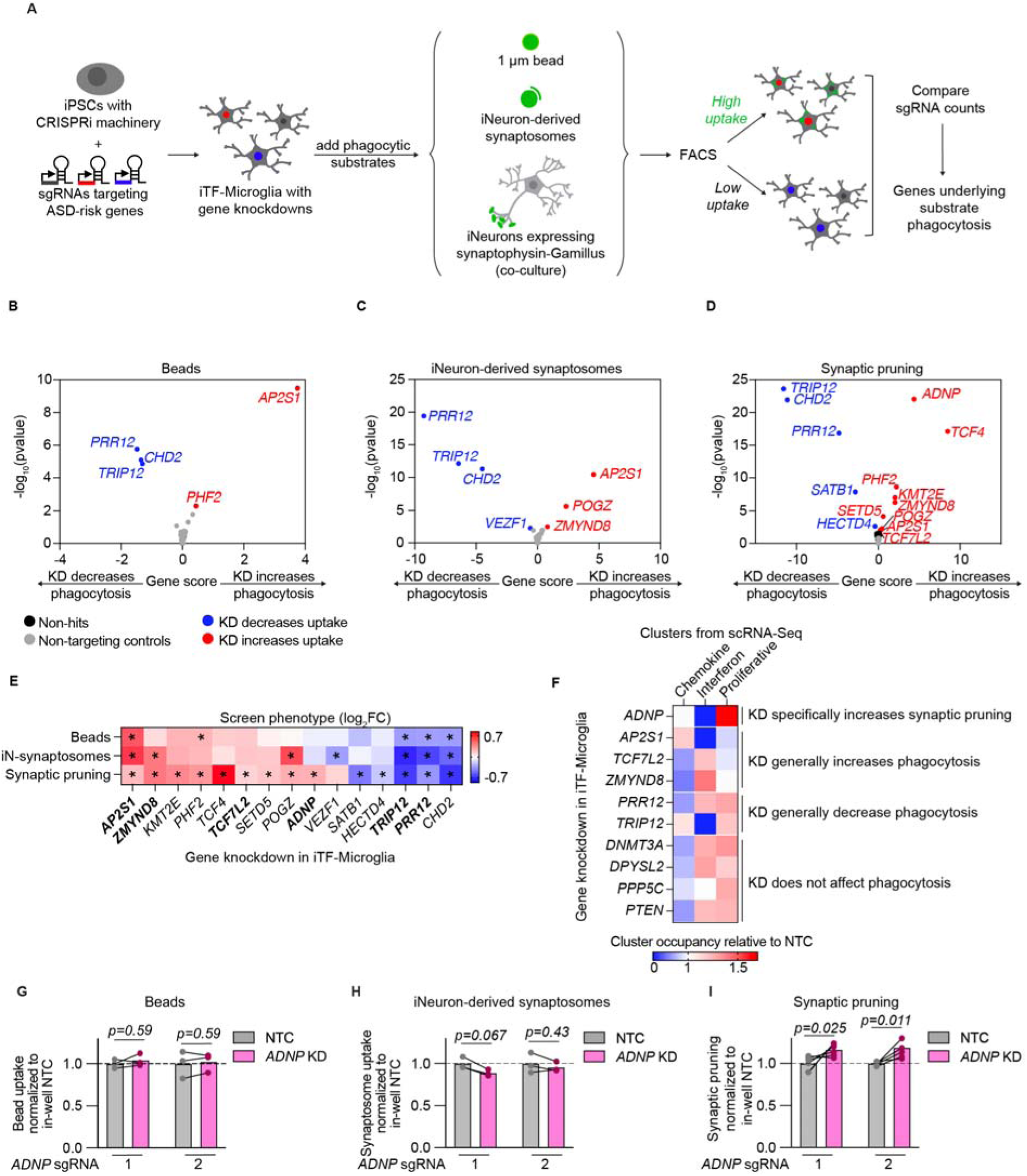
Convergent and divergent phagocytosis phenotypes reveal *ADNP* as a unique genetic regulator of iPSC-derived microglia (iTF-Microglia). A. CRISPRi-based screening strategy to assess the role of ASD risk genes in microglial phagocytosis of various substrates: 1 um beads, iNeuron-derived synaptosomes, and live synaptic material. B-D. Volcano plots showing the effect of gene knockdowns (KDs) in iTF-Microglia on uptake of beads (B), iNeuron-derived synaptosomes (C), and live synapses (D) across three screening replicates and their corresponding statistical significance (two-sided Mann–Whitney U-test, false discovery rate (FDR = 0.01). Gene KDs that increase uptake are noted in red, gene KDs that decrease uptake are noted in blue, non-targeting controls (NTCs) are noted in gray, and non-hit gene knockdowns are noted in black. E. Heatmap comparing the phagocytosis phenotypes across substrates for each hit gene. Statistically significant phenotypes (FDR < 0.01) are noted with an asterisks. Bolded gene KDs are further analyzed in F. F. Heatmap comparing the cluster occupancy of cells with a given gene KD relative to NTCs. Increased cluster occupancy is noted by reds, and decreased cluster occupancy is noted by blues. Gene KDs are grouped by their phagocytosis phenotypes. G-H. Individual validation of phagocytosis phenotypes for iTF-Microglia with *ADNP* KD (N = 6 wells, multiple paired t-tests).

Additionally, screen hits showed consistent phenotypes across their two sgRNAs, with the exception of one *CHD2* targeting sgRNAs which showed divergent phenotypes as well as no knockdown (Supplementary Figure 4E,F). Aggregating phagocytosis phenotypes across screens for all gene knockdowns that were significant in at least one screen, we see that many gene knockdowns had similar effects across substrates (Figure 3E). Notably, *ADNP* knockdown specifically increased iTF-Microglia uptake of live synaptic material. We further assessed points of convergence and divergence for ASD risk genes by comparing the transcriptomic activation states to the phagocytosis phenotypes induced by specific ASD risk gene knockdowns (Figure 3F). Again notably, *ADNP* KD showed a stark depletion of the interferon activation state and enrichment of the proliferation activation state when compared to other gene knockdowns and phagocytosis phenotypes.

Previous work has shown mice with haploinsufficient Adnp have decreased spine density and developmental delays^40–42^. In neurons, *ADNP* plays a role in processes including differentiation^43,44^, microtubule stability^45,46^, and neurite outgrowth^47^ but little is known about the role ADNP plays in microglia and how it may affect uptake of synaptic material and activation state. Towards this end, we first validated the screen phenotypes for individual sgRNAs targeting *ADNP* (Figure 3G-I, Supplementary Figure 4G). We next asked why iTF-Microglia with *ADNP* knockdown specifically increased uptake of live synapses.

### *ADNP* knockdown (KD) alters iPSC-derived microglia (iTF-Microglia) proteome and interactions with iPSC-derived neurons (iNeurons)

Microglial interactions with neurons are dependent on surface expression of many receptors. To assess differential abundance at the protein level, we performed surface proteomics as well as whole cell proteomics for iTF-Microglia with *ADNP* knockdown versus an NTC sgRNA (Figure 4A,B). Both datasets revealed significantly differentially abundant proteins (DAPs) important for cell motility (SN, CD163, NUMA1, FINC), endocytosis (RAB18, AP2S1, RAB35, RAC1), and synaptic pruning (C1QC, ANXA2, ANXA4, ANXA6). Comparing the changes induced by *ADNP* loss at the surface and in the whole cell (Figure 4C), we saw a general correlation between surface protein changes and whole cell changes. However, some proteins that changed at the surface did not change at the whole-cell level, suggesting defects in receptor translocation and recycling. Indeed, GO Cellular Component enrichment identified broad changes in intracellular vesicle and cytoskeletal components (Figure 4D).

**Figure 4.**
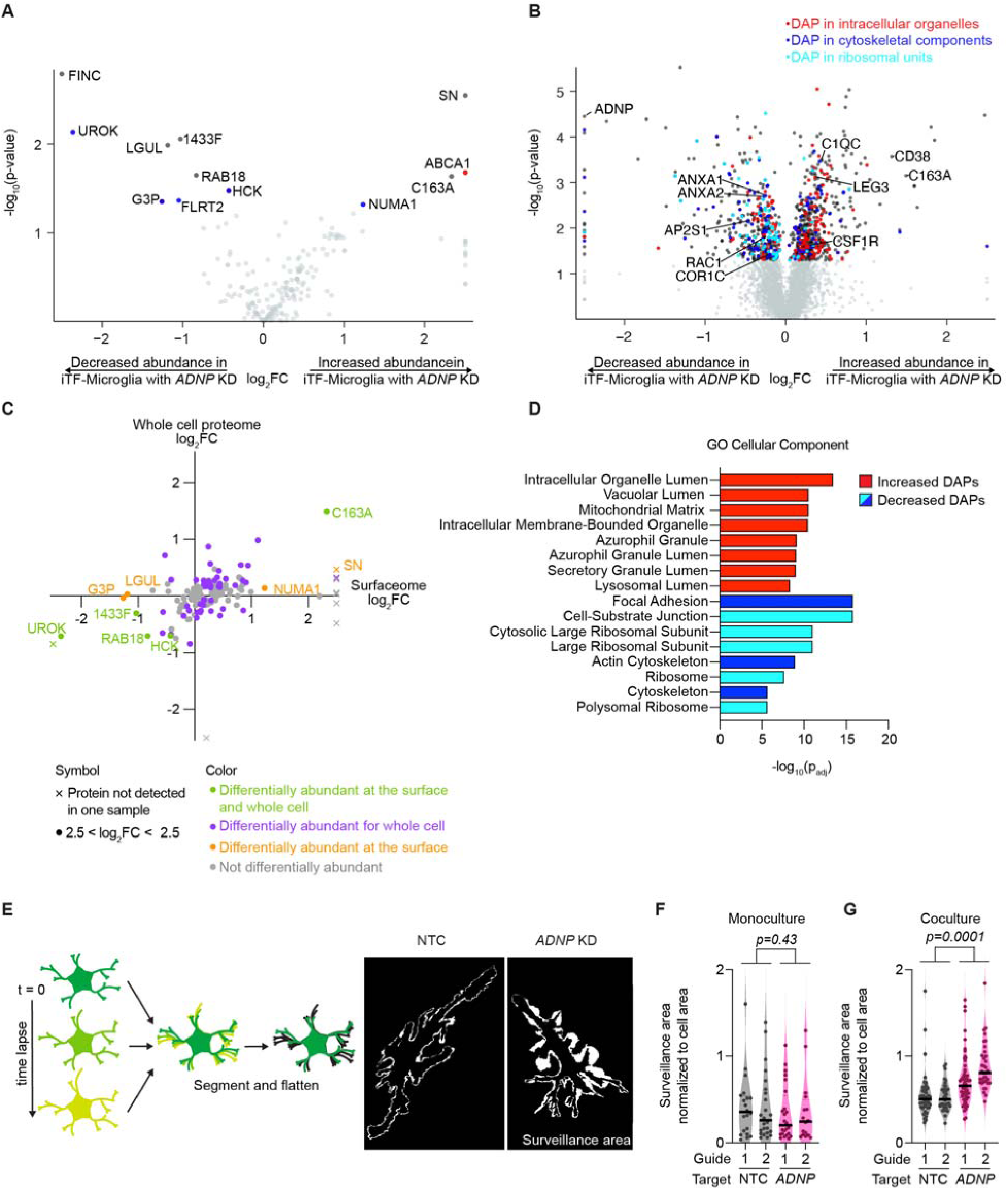
*ADNP* knockdown (KD) alters iPSC-derived microglia (iTF-Microglia) proteome and interactions with iPSC-derived neurons (iNeurons). A-B. Volcano plots showing the change in abundance for individual surface proteins (A) or whole cell proteins (B) in iTF-Microglia with *ADNP* KD compared to iTF-Microglia with non-targeting control (NTC) sgRNA as determined by surface protein labeling and mass spectroscopy (N = 3 replicates per cell type). Proteins with statistically significant changes in abundance in iTF-Microglia with *ADNP* KD are noted in dark gray and those contributing to specific GO Cellular Component pathways in D are correspondingly colored red, blue, or cyan. Proteins with no statistically significant change are noted in light gray (Differentially expressed proteins (DAPs), Padjl<l0.05, two-tailed Student’s t-test). Genes whose expression turned on or off with *ADNP* KD are marked by an X symbol. C. Scatterplot comparing protein changes at the surface and whole cell for iTF-Microglia with *ADNP* KD. Proteins that are significantly differentially expressed in both datasets are noted in green, those exclusive to the whole cell are noted in purple, those exclusive to the cell surface are noted in orange, and those with no significant differential expression are noted in gray. D. GO Cellular Component enrichment analyses of DAPs with increased (red) or decreased (blues) expression in iTF-Microglia with *ADNP* KD (Padj < 0.05). Terms are color-coded such that terms of the same color have many overlapping genes. E. (Left) Surveillance area quantification strategy. Time lapse immunofluorescence micrographs of iTF-Microglia expressing a fluorescently tagged membrane protein, Lck-mNeonGreen, are acquired, segmented, and maximum-intensity projections are analyzed to measure surveillance area. (Right) Representative surveillance areas from coculture iTF-Microglia with *ADNP* KD or NTC. F. Single cell surveillance area normalized to initial cell area for monoculture iTF-Microglia with *ADNP* KD or NTC (N = 16-24 single cells, violin plot area represents the density curve, significance based on NTC vs ADNP factor computed by two-way ANOVA). G. Single cell surveillance area normalized to initial cell area for coculture of iTF-Microglia with *ADNP* KD or NTC with iNeurons (N = 35-59 single cells, violin plot area represents the density curve, t significance based on NTC vs ADNP factor computed by two-way ANOVA).

Together, the changes in proteins related to motility and the cytoskeleton with *ADNP* loss led us to ask whether iTF-Microglia with *ADNP* knockdown altered microglial motility and surveillance dynamics. We developed a live cell imaging pipeline in which we used membrane labeled iTF-Microglia, acquired time lapses, segmented individual microglia, and compared the total area surveilled by the microglia to its initial area (Figure 4E). Comparing iTF-Microglia with *ADNP* knockdown to iTF-Microglia with NTC in monoculture, we saw no difference in surveillance area (Figure 4F). Interestingly, comparing the microglia in coculture, we saw that iTF-Microglia with *ADNP* knockdown had a larger surveillance area than iTF-Microglia with NTC (Figure 4G). The difference in surveillance area was not driven by differences in microglia morphology in monoculture or coculture (Supplementary Figure 5A-H). We questioned if increased motility in coculture explained why we only saw phagocytic differences with synaptic pruning, but *ADNP* knockdown still showed no change in synaptosomes phagocytosis even in coculture (Supplementary Figure 5I), suggesting that the remodeled proteome and corresponding motility changes of iTF-Microglia with *ADNP* loss specifically affects microglia interactions with live neurons and synapses.

### *ADNP* loss affects endocytic trafficking in iPSC-derived microglia (iTF-Microglia)

To evaluate if proteomic changes in iTF-Microglia with *ADNP* knockdown are driven by transcriptional changes, we investigated the differentially expressed genes (DEGs) of iTF-Microglia with *ADNP* knockdown compared to iTF-Microglia with NTC (Figure 5A). GO Cellular Component enrichment identified increased expression of genes relating to endolysosomal machinery (Figure 5B). Enrichment analysis of downregulated DEGs revealed no statistically significant gene sets (Supplemental Figure 6A). The endolysosomal signature is distinct from the cytoskeletal changes suggested by DAPs. This difference was further observed when directly comparing all changes in mRNA and corresponding protein expression with *ADNP* loss (Figure 5C). This direct comparison revealed little correlation between mRNA and protein levels, an observation reported in other microglia studies^48–50^. Investigating the specific DEGs and DAPs, we also found very little overlap (Figure 5D), suggesting that changes in the cell surfaceome and proteome were due to a combination of transcriptional and post-transcriptional mechanisms.

**Figure 5:**
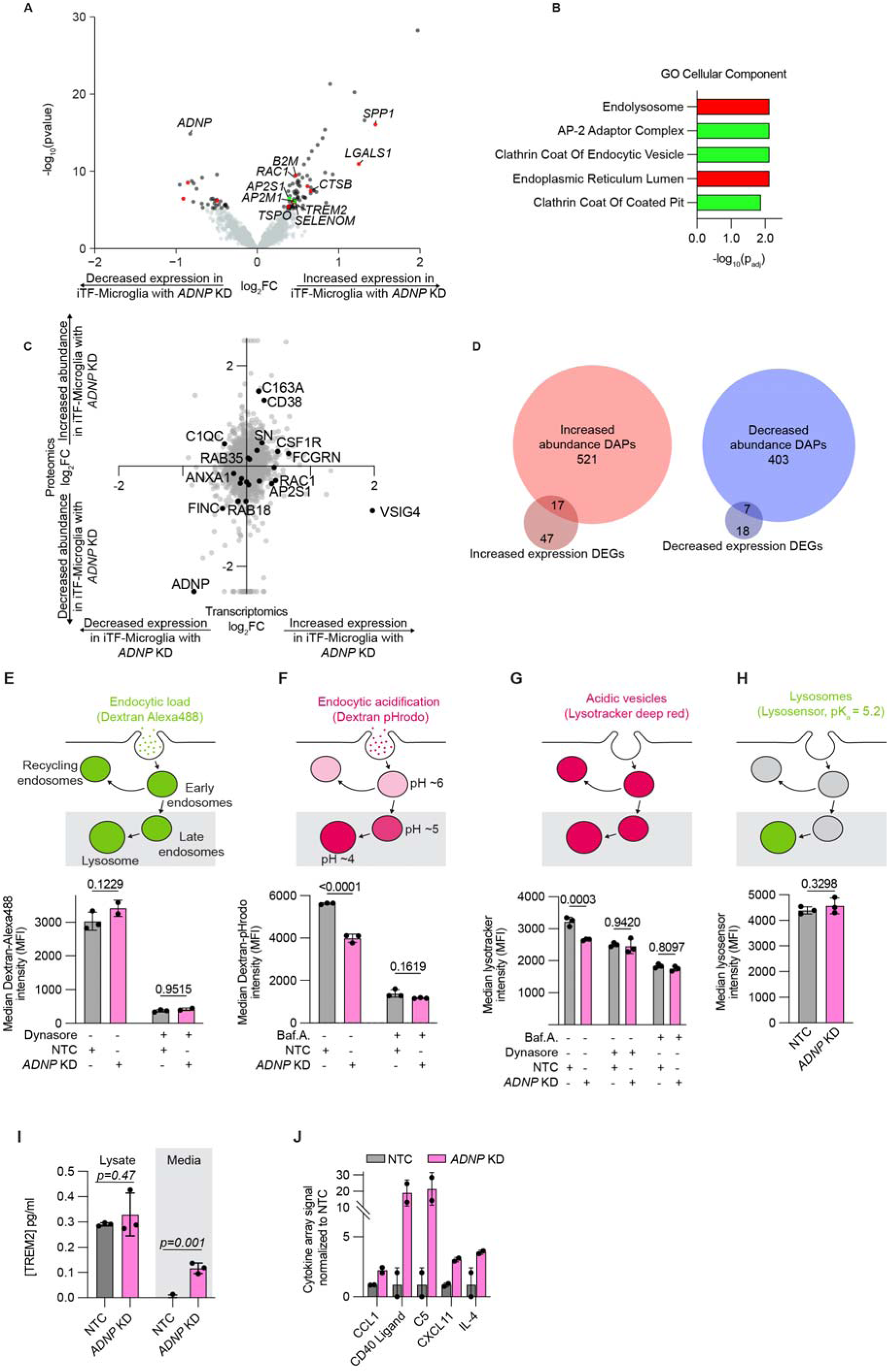
*ADNP* knockdown (KD) affects endocytic trafficking in iPSC-derived microglia (iTF-Microglia). A. Volcano plot showing the change in expression as measured by RNA-sequencing for individual genes in iTF-Microglia with *ADNP* KD compared to iTF-Microglia with non-targeting control (NTC) sgRNAs. RNA-sequencing data extracted from a CROP-seq dataset, see Methods for detail. Significantly differentially expressed genes (DEGs) are noted in dark gray or color-coded based on their corresponding categorization in enrichment analyses (B) and genes with no statistically significant change are noted in light gray (DEGs, Padjl<l0.05, two-tailed Student’s t-test). B. GO Cellular Component enrichment analyses of DEGs with increased expression in iTF-Microglia with *ADNP* KD. Terms representing unique gene sets and with Padj < 0.05 are displayed. Terms are color-coded such that terms of the same color have many overlapping genes. C. Scatterplot comparing fold changes at the transcriptomic and proteomic levels when comparing iTF-Microglia with *ADNP* KD to iTF-Microglia with NTC. Some significantly differentially abundant proteins are noted in black and labeled. D. Venn diagrams showing the overlap in differentially expressed genes and proteins that are increased or decreased in iTF-Microglia with *ADNP* KD. E. (Top) Schematic of the biology of dextran-Alexa488 endocytosis. (Bottom) Endocytic load within iTF-Microglia with *ADNP* KD or NTC and with or without endocytosis inhibitor, Dynasore, as measured by flow cytometry (N = 2-3 wells, bars represent mean +/-standard deviation, 2 way ANOVA). F. (Top) Schematic of the biology of dextran-pHrodo endocytosis. (Bottom) Endocytic acidification within iTF-Microglia with *ADNP* KD or NTC and with or without V-ATPase inhibitor, BafilomycinA, as measured by flow cytometry (N = 3 wells, bars represent mean +/- standard deviation, 2 way ANOVA). G. (Top) Schematic of the biology of Lysotracker. (Bottom) Acidic vesicles within iTF-Microglia with *ADNP* KD or NTC and with or without Dynasore and BafilomycinA as measured by flow cytometry (N = 3 wells, bars represent mean +/- standard deviation, 2 way ANOVA). H. (Top) Schematic of the biology of lysosensor. (Bottom) Lysosomes within iTF-Microglia with *ADNP* KD or NTC as measured by flow cytometry (N = 3 wells, bars represent mean +/- standard deviation, two-tailed Student’s t-test). I. TREM2 concentration of cell lysates or media collected from iTF-Microglia with *ADNP* KD or NTC measured using a Homogeneous Time Resolved Fluorescence (HTRF) assay (N = 3 wells, bars represent mean +/- standard deviation, t-test). J. Relative abundances measured using the integrated intensity from a cytokine array for proteins that have a trend for differential abundance between iTF-Microglia with NTC or *ADNP* KD (N = 2 wells, bars represent mean +/- standard deviation).

One such mechanism, suggested by the transcriptomic data, is receptor endocytosis and protein trafficking. We first assessed endocytic load by incubating iTF-Microglia with Alexa488 labeled dextran (Figure 5E). Comparing iTF-Microglia with NTC or *ADNP* knockdown, we saw no difference in endocytic load. Treatment with Dynasore, a dynamin-mediated endocytosis inhibitor, also showed no difference in Dextran-Alexa488 uptake. An important feature of trafficking – to the lysosome, Golgi complex, or back to the cell surface – after endocytosis is vesicle acidification. To assess endocytic acidification, we incubated iTF-Microglia with pHrodo (a pH sensitive dye) labeled dextran (Figure 5F). As endocytic vesicles become acidified, the pHrodo fluorescence intensity increases. Comparing iTF-Microglia with NTC or *ADNP* knockdown, we saw a decrease in endocytic acidification. Treatment with Bafilomycin A, a V-ATPase and acidification inhibitor, showed no difference in Dextran-pHrodo uptake. Using an orthogonal dye, Lysotracker, to measure acidic vesicles in cells, we also observed a decrease in acidic vesicles in iTF-Microglia with *ADNP* knockdown compared to iTF-Microglia with NTC (Figure 5G). This difference was abolished by treating cells with Bafilomycin A as well as Dynasore, increasing our confidence that the difference in acidic vesicles might be downstream of endocytosis. Further, we probed lysosome content using Lysosensor, and did not observe a difference between iTF-Microglia with NTC and *ADNP* knockdown (Figure 5H). All flow cytometry observations were also recapitulated by microscopy (Supplemental Figure 6B-E). Additional analysis of lysosomal protease activity did not reveal differences between iTF-Microglia with NTC and *ADNP* knockdown except at long time points, at which iTF-Microglia with *ADNP* knockdown show a slight increase in protease activity (Supplemental Figure 6F).

The transcriptomic data also showed an increase in *TREM2* expression in iTF-Microglia with ADNP knockdown compared to iTF-Microglia with NTC. TREM2 is an important receptor for microglial activation and uptake of synaptic pruning. Using a Homogeneous Time Resolved Fluorescence (HTRF) TREM2 assay, we did not find an increase in TREM2 protein in cell lysates, but we did observe an increase in TREM2 in the culture supernatant (Figure 5I). TREM2 release from microglia is hypothesized to reflect activated TREM2 signaling^51^. This observation of increased extracellular TREM2 led us to ask if other secreted proteins are differentially regulated by *ADNP* loss. We used an arrayed cytokine immunoassay to assess the secretome of iTF-Microglia with *ADNP* knockdown compared to iTF-Microglia with NTC (Figure 5J and Supplementary Table 3). iTF-Microglia with *ADNP* knockdown had elevated secretion of CCL1, CD40 ligand, C5, CXCL11, and IL-4. Elevated C5 and IL-4 have been found to be associated with ASD in patients^52,53^. The changes in these protein levels was not driven by changes at the transcriptomic level (Supplementary Figure 6G). Taken together, we observed changes in endocytic vesicle trafficking, surface proteome, and secretome of iTF-Microglia with ADNP loss.

### ADNP localizes to early endocytic compartments in iPSC-derived microglia (iTF-microglia)

Our characterization of *ADNP* loss in iTF-Microglia highlights distinct cell phenotypes when compared to previous work studying the role of *ADNP* in neurons. Additionally, when we investigated the localization of ADNP in iTF-Microglia compared to iNeurons, we saw a stark difference (Figure 6A). Consistent with previous work, ADNP was localized in the nucleus and cytoplasm^54,55^ of our iNeurons with about 90% of neurons positive for nuclear ADNP (Figure 6B). The localization of ADNP in iTF-Microglia differed from that of neurons, showing a strong extranuclear punctate pattern and a low percentage of microglia positive for nuclear ADNP. The localization difference persisted when iNeurons and iTF-Microglia were cocultured (Supplementary Figure 7A-C).

**Figure 6:**
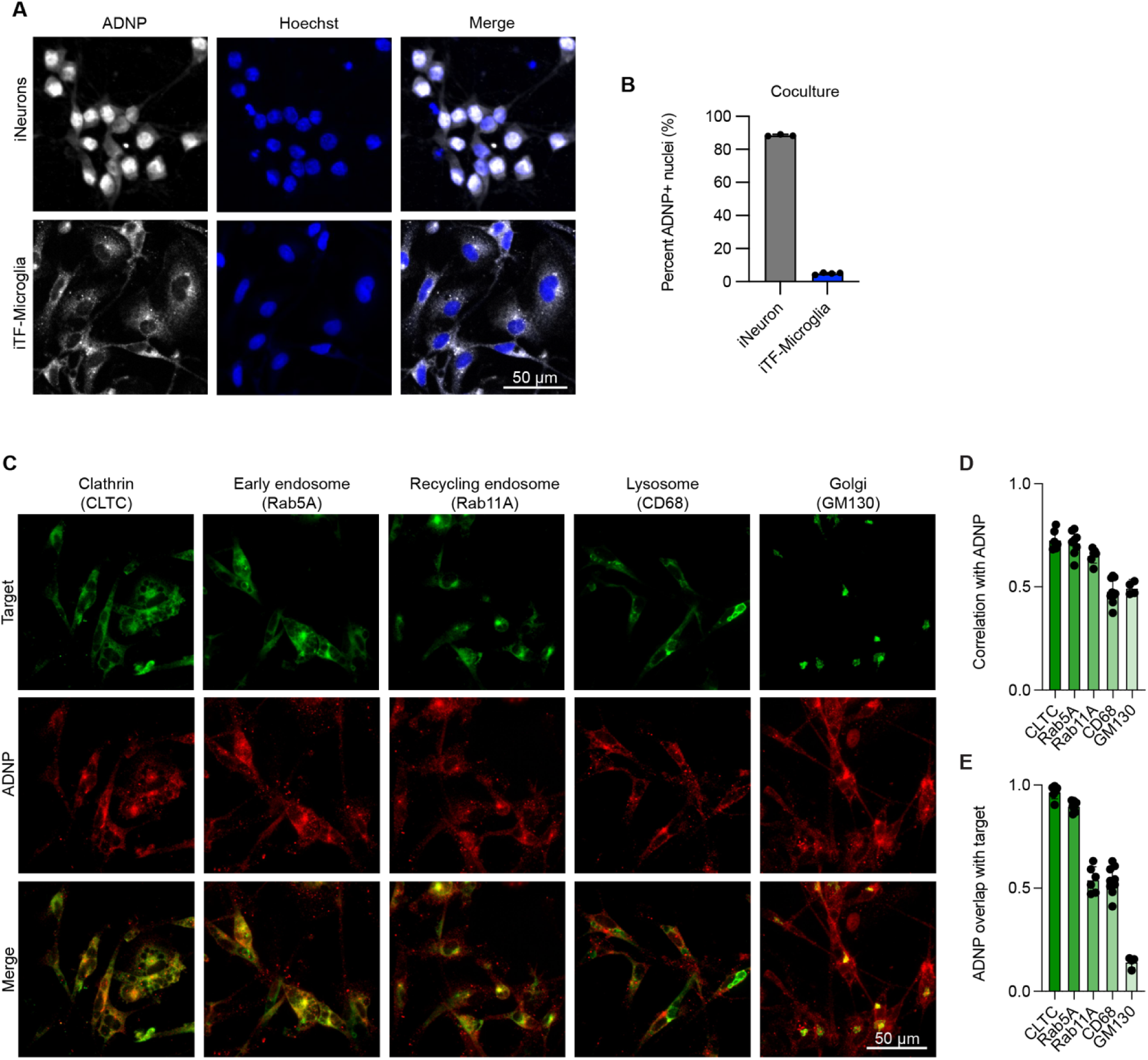
ADNP localizes to early endocytic compartments in iPSC-derived microglia (iTF-microglia). A. Representative micrographs of iPSC-derived neurons (iNeurons) and iTF-Microglia showing the localization of ADNP (detected by immunofluorescence, gray) with respect to nuclei demarked by Hoechst 33342 (blue). B. Quantification of the percent of nuclei with high ADNP intensity (N = 3-4 wells per cell type, bars represent mean +/- standard deviation). C. Representative immunofluorescence micrographs of iTF-Microglia stained with antibodies against ADNP (red) and various endolysosomal organelle markers including CLTC, Rab5A, Rab11A, CD68, and GM130 (green). D. Average Pearson correlation between ADNP and individual organelle markers measured per cell for a given field of view (N = 1-4 fields of view from 3 wells, bars represent mean +/- standard deviation). E. Average Mander’s overlap for ADNP and individual organelle markers measured per cell for a given field of view (N = 1-4 fields of view from 3 wells, bars represent mean +/- standard deviation).

Based on the localization pattern in iTF-Microglia, we hypothesized the ADNP was localizing to endosomal compartments. We co-stained iTF-Microglia for ADNP and markers of various endosomal structures including clathrin, early endosomes, recycling endosomes, lysosomes, and the Golgi apparatus (Figure 6C). Using correlation and overlap analyses, we found that much of ADNP co-localized with clathrin and early endosome markers, although there was overlap with the other endosomal structures as well (Figure 6D,E). Altogether, our transcriptomic, localization, and functional assays suggest that *ADNP* loss affects endocytosis in iTF-microglia, possibly due to a direct role that ADNP plays in this process.

### Altered endocytosis is seen in human patient data

To assess if the pathways we have identified in our iPSC-derived models are relevant to patient phenotypes, we performed pathway enrichment analysis for upregulated DEGs from primary human microglia from postmortem samples from individuals with ASD and neurotypical controls^56^. The most significant GO Cellular Component pathways upregulated in this dataset point to ribosomal subunits, nucleus, and, interestingly, focal adhesion and endosomal compartments (Figure 7A,B and Supplementary Table 2). The conservation of upregulated endosomal compartments between these two datasets suggests that changes in endosomal pathways in microglia may be a common feature in ASD patients, beyond those with *ADNP* loss of function. Altogether, we report the *ADNP* loss results in altered endocytic trafficking, a remodeled proteome, increased motility, and a specific increase in synaptic pruning. We hypothesize that altered endocytic trafficking affects receptor recycling that is important for both motility and synaptic pruning (Figure 7C).

**Figure 7:**
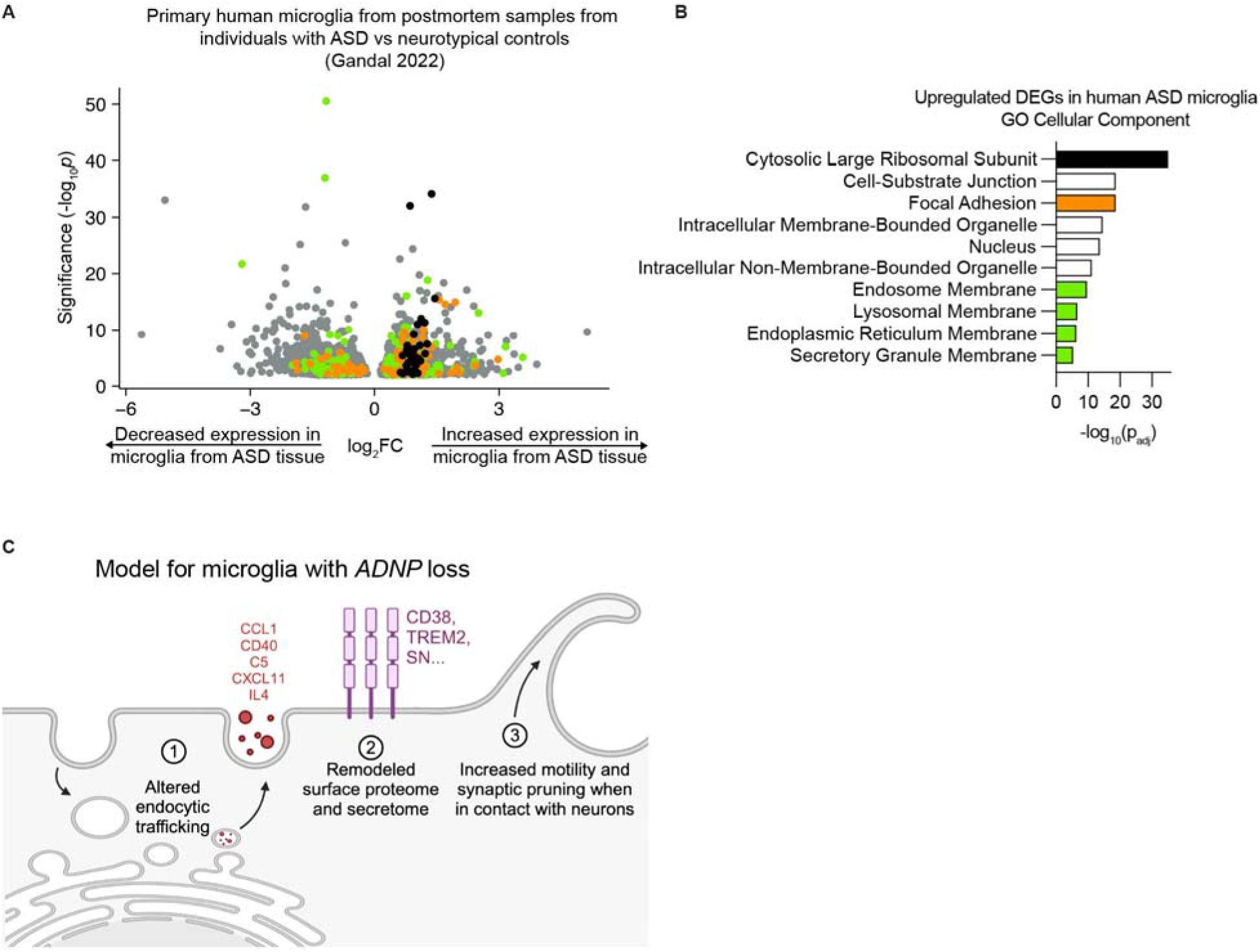
Altered endocytosis in microglia is supported by transcriptomic data from human ASD brain tissue. A. Volcano plot showing the change in expression as measured by single cell RNA-sequencing for differentially expressed genes (DEGs) in microglia from postmortem tissue from individuals with ASD compared to neurotypical controls^37^ (DEGs, Benjamini & Hochberg corrected p values < 0.01). B. GO Cellular Component enrichment analysis of DEGs with increased expression in microglia from postmortem tissue from individuals with ASD. Terms representing unique gene sets and with Padj < 0.05 are displayed. Terms are color-coded such that terms of the same color have many overlapping genes. C. Summary model of microglial phenotypes with *ADNP* loss.

## DISCUSSION

In this study, we developed an iPSC-derived microglia-neuron coculture and flow cytometry method for measuring synaptic pruning, which enabled us to conduct parallel CRISPR-based phagocytosis screens on beads, synaptosomes, and synaptic pruning to systematically assess the effect of ASD-risk genes on microglial phagocytosis. Recently published work presented a similar flow cytometry method for measuring microglial uptake of Alexa Fluor-labeled retinal ganglion cell (RGC) material in the lateral geniculate nucleus (LGN) after optic nerve crush^57^. Synapse remodeling of RGC inputs in the LGN is well defined such that internalization of Alexa Fluor-labeled material by microglia in the LGN can be attributed to synaptic pruning^13^. Since synaptic pruning is less understood *in vitro*, we designed the synaptophysin-Gamillus construct to provide confer higher specificity of synaptic material compared to general neuronal material. Compared to imaging readouts of synaptic pruning, our flow cytometry readout does not remove background signal. With imaging methods, synaptic segmentation algorithms remove the contribution of background fluorescence. In our readout, measuring total fluorescence of the cell by flow cytometry means we include background autofluorescence. This methodological difference leads to smaller effect sizes, but the high throughput and sensitivity of flow cytometry increases the robustness in relative differences of synaptic pruning. This engineered platform can be used to assess synaptic pruning with other genetic perturbations, such as knockdown of genes associated with other neurodevelopmental or neurodegenerative disorders, in microglia and neurons or pharmacological perturbations. Importantly, this *in vitro* system can facilitate the molecular interrogation of synaptic pruning mechanisms.

The combination of transcriptomic profiles from single cell sequencing and phagocytosis phenotypes from the three screens allowed us to investigate points of mechanistic convergence by which ASD risk genes may act in microglia. Surprisingly, we did not find transcriptomic changes in genes known to mediate phagocytosis. However, we did find an anti-correlation between interferon and chemokine activation states. ASD risk gene knockdowns that increased occupancy in the interferon state showed a decrease in the chemokine state, a relationship others have reported in various contexts^27,58,59^. Increased interferon response has been previously reported in human ASD transcriptomic datasets^7,56^. This interferon and chemokine signature may represent one point of convergence describing a microglial state in ASD, but the functional impact of this pro-interferon state may not be robustly affecting phagocytosis.

From those ASD risk genes that do affect microglial phagocytosis, *ADNP* emerged as a notable outlier that specifically affected synaptic pruning. Through transcriptomic, proteomic, and functional characterization of microglia with *ADNP* knockdown, we uncovered that loss of *ADNP* in microglia resulted in decreased endocytic trafficking, remodeled proteome, and increased motility of microglia cocultured with neurons. We observed a multifaceted proteomic remodeling of many components implicated in synaptic pruning (decreased annexins^38^, increased C1QC^13^, increased TREM2^37^) and motility (CD163, FCGRN, SN, ABCA1, NUMA1, FINC). By comparing phagocytosis of synaptosomes in monoculture to synaptic pruning and motility in monoculture and coculture, we see that iTF-Microglia with *ADNP* knockdown specifically affects these phenotypes in coculture, suggesting that the proteomic remodeling induced by *ADNP* loss affects microglia-neuron interactions. Transcriptomic changes largely did not explain proteomic changes which others have also reported in microglia^48–50^. We hypothesize that many proteomic changes in iTF-Microglia with *ADNP* knockdown are explained by deficits in endocytic trafficking that result in abnormal receptor recycling and degradation.

Microglial activation at the transcriptomic level has previously been reported in a study using shRNA-mediated knockdown of *ADNP* with AAV9 delivery in mice^42^. AAV9 selectively infects neural cells, so subsequent microglia activation is induced in a cell non-autonomous manner. Microglia activation in response to *ADNP* knockdown in neural cells included increased microglia density, increased CD68 staining, and increased complement components. In contrast to that work, we provide direct evidence that *ADNP* knockdown in human microglia causes increased synaptic pruning. It is complementary to their transcriptomic evidence of microglia activation upon *ADNP* knockdown in mouse neural cells, but distinct in that our evidence is causal as well as functional. Interestingly, the AAV9 delivery of shRNA targeting *ADNP* also uncovered overlapping microglia and astrocyte differentially expressed genes. Although we were not able to test *ADNP* knockdown in astrocytes, it is possible similar phenotypes could be seen in astrocytes. With respect to neurons, previous work from our group showed that *ADNP* knockdown in human iPSC-derived neurons (iNeurons) decreases survival^60^. Therefore, knockdown of *ADNP* in iNeurons may have more pleiotropic effects that would confound interpretations of microglia-neuron interactions in our model system.

From our work and others, we see that ADNP is expressed more highly in neurons than in microglia, and it is thus likely that a large contribution to autism phenotypes is due to neuronal dysfunction; however, microglia dysregulation is commonly seen in ASD patient brain tissue^4–8^, and mechanisms regulated by low level expression of ASD risk proteins in microglia could contribute. Our work supports this idea and adds to our understanding of the biology of ADNP syndrome by investigating the specific effect of *ADNP* loss in microglia. We offer evidence that *ADNP* and other ASD risk genes can induce an activated microglia state in a cell-autonomous manner.

The cell-type specific localization of ADNP, nuclear in iNeurons and cytoplasmic in iTF-Microglia, reveals dynamic regulation of this protein. Others have reported the cytoplasmic localization of ADNP mediated by 14-3-3 proteins to facilitate neurite extension^44,55^ and abnormal cytoplasmic localization of ADNP induced by patient mutations^61,62^, suggesting that the localization of ADNP is important for its function. Interestingly, our proteomics reveal the decreased expression of 14-3-3 F in iTF-Microglia with *ADNP* knockdown. This may be explained by a shared coexpression network and interaction between these two proteins in our iTF-Microglia.

Our study has several limitations. One is the use of iPSC-derived microglia. iPSC-derived microglia are known to be basally more activated than those cells found in tissue^27,63^. The agreement in transcriptomic signatures between our iTF-Microglia with *ADNP* knockdown and those found upon shRNA-mediated *ADNP* knockdown^42^ in mice suggests that we are still able to detect meaningful activation states. Second, the physiological relevance of synaptic pruning in an iPSC-derived microglia-neuron coculture is not known. Many studies of synaptic pruning in *in vivo* mouse models investigate well characterized periods of activity-dependent synaptic remodeling, like eye input segregation in the visual cortex^13^. However, our system reproduces key aspects conserved in *in vivo* assays, such as decreased uptake of synaptic material by blocking phosphatidylserine^38,39^, suggesting principles of synaptic pruning may hold true in our experimental paradigm. Lastly, the CROP-Seq dataset is limited by low cell counts such that we were not able to assess more than 12 ASD risk gene knockdowns.

In summary, we used CRISPRi-based screens to systematically assess transcriptomic activation and phagocytic phenotypes of iPSC-derived microglia with ASD risk gene knockdown. This functional genomics approach allowed us to nominate ASD risk genes that affect activation and phagocytosis. We did not observe many global trends of activation or phagocytosis, suggesting that these phenotypes may not be robust points of convergence by which many ASD risk genes act. Our work does support the need for further study of ASD risk genes in microglia and in microglia-neuron interactions.

## Data availability

All datasets will be made available at NCBIGeo and CRIPSRbrain (https://crisprbrain.org/) at the time of publication.

## Code availability

All code, including R notebooks for transcriptomic analyses and CellProfiler pipelines for image analyses, will be made available at https://kampmannlab.ucsf.edu/resources at the time of publication.

## Acknowledgements

We thank Illana Gozes for advice and insights into studying *ADNP*, Caroline Mrejen and the Innovation Core at the Weill Institute for Neurosciences for sharing their microscopy expertise, Sarah Elmes for sharing her flow cytometry and FACS expertise. We thank Nawei Sun and Jeremy Willsey for sharing sgRNA oligos. We acknowledge and thank all members of the Kampmann lab for their technical support and positive scientific community. This work was supported by the National Science Foundation Graduate Research Fellowship under Grant No. 2034836 (to O.T.), NIH/NIMH grant U01 MH115747 (to M.K.), Alzheimer’s Association grant AARF-22-973222 (to A.M.), Larry L. Hillblom Foundation frant 2022-A-016-FEL (to A.M.), R01 MH125516 (to T.J.N.)

## Author Contributions

The CRISPRi-based screen was designed, conducted, and analyzed by OT and MK. Significant intellectual contributions from AM, VH, WL. The CROP-Sequencing experiment was conducted and analyzed by ND, SS, and OT. ADNP follow up experiments were conducted by OT with BH leading the design, experiment, and analysis of surface proteomics and AM contributing to the TREM2 experiment design and analysis. Molecular cloning conducted by OT with help from VCC, VP, KL, and SB. OT and MK created the figures and wrote the manuscript with contributions from AM. All authors reviewed and approved the final manuscript.

## Competing Interests

M.K. is a co-scientific founder of Montara Therapeutics and serves on the Scientific Advisory Boards of Montara Therapeutics, Engine Biosciences, Casma Therapeutics, Alector, and Neurocrine, and is an advisor to Modulo Bio and Recursion Therapeutics. M.K. is an inventor on US Patent 11,254,933 related to CRISPRi and CRISPRa screening, and on a US Patent application on *in vivo* screening methods.

---

**Supplemental Fig. 1:**
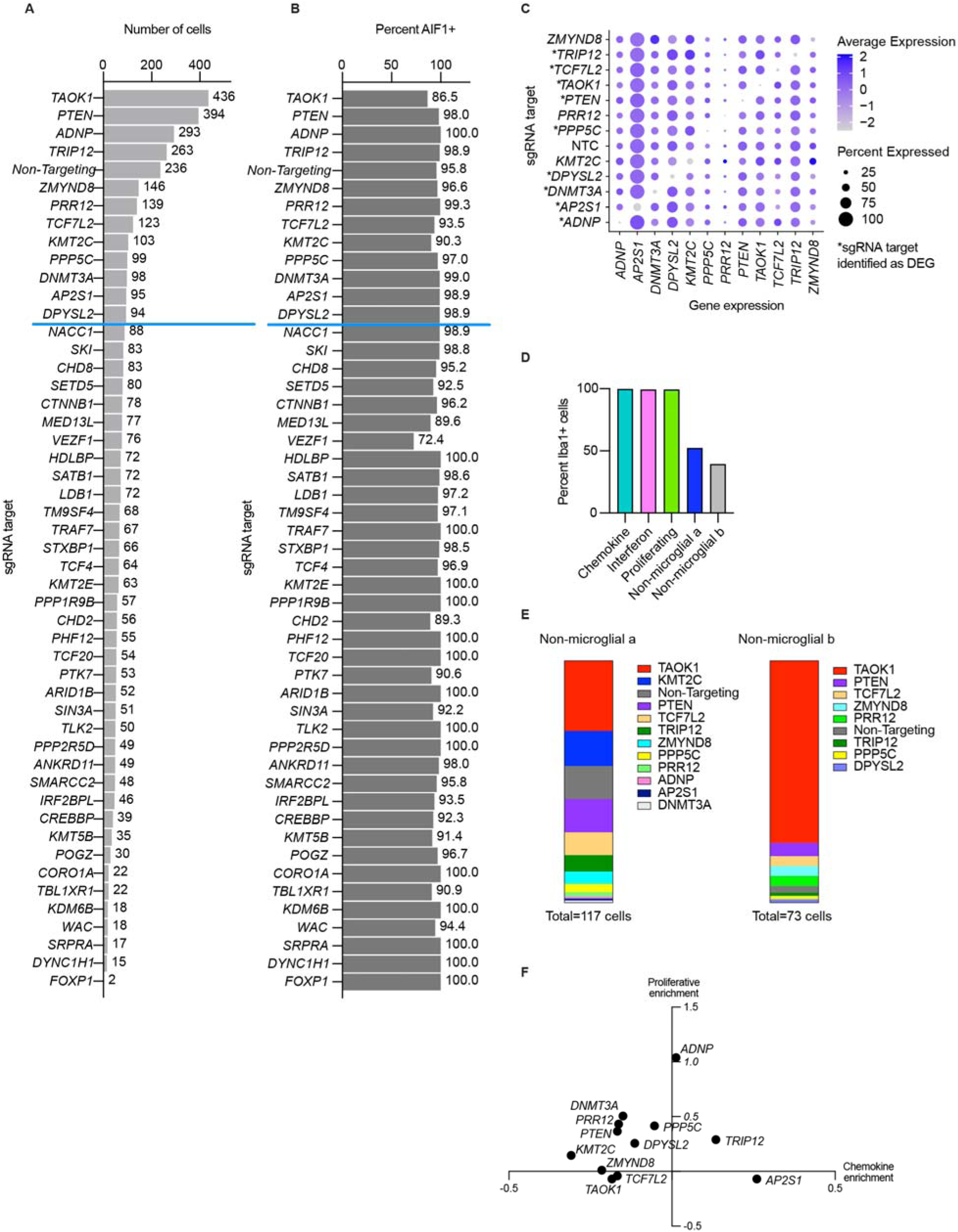
CROP-Seq quality control. A. Bar graph of the number of cells for each gene knockdown (KD). Gene KDs with greater than 90 cells were further studied. B. Bar graph of the percent Iba1 or *AIF1* positive cells for each gene KD. C. Dot plot displaying the efficacy of gene KDs (x axis) across 12 different gene KDs and non-targeting control sgRNAs (y axis). The color of the dots reflect the relative expression of the gene, and the size of the dots represent the percent of cells expressing the gene for a given gene KD. D. Percent *AIF1* (Iba1) positive cells in each cluster. E. Composition of non-microglial clusters a and b in terms of cells and their specific gene knockdown. F. Scatterplot showing the change in the proliferative and chemokine clusters relative to non-targeting controls.

**Supplemental Fig. 2:**
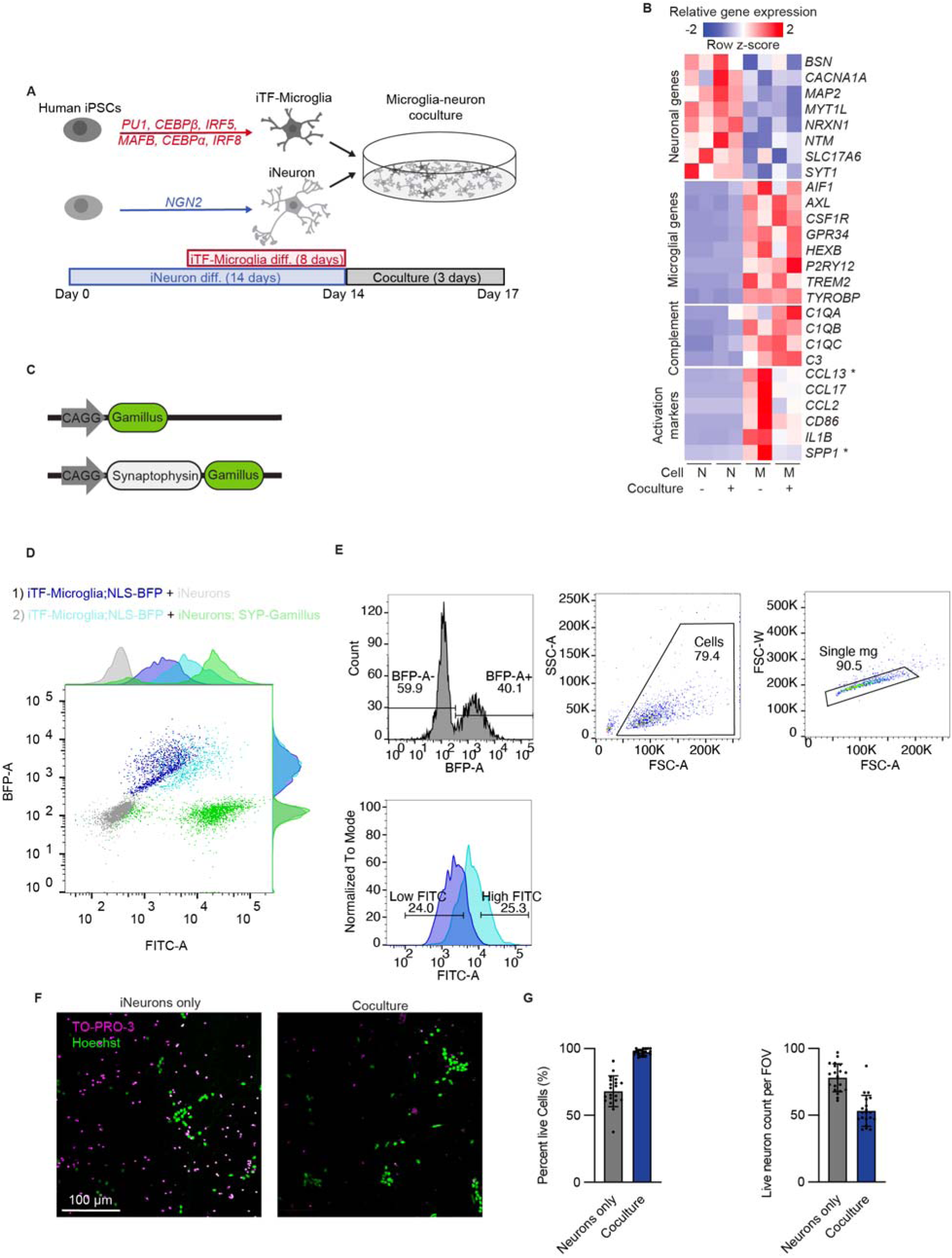
Characterization of cocultures of iPSC-derived neuron (iNeuron) and iPSC-derived microglia (iTF-Microglia) and synaptic material uptake. A. iTF-Microglia and iNeuron differentiation and coculture strategy. B. Heatmap showing relative gene expression from RNA sequencing of neuronal genes, microglial genes, complement components, and microglial activation markers by iTF-Microglia (M) and iNeurons (N) in monoculture and coculture preparations (N = 2 biological replicates). C. Gamillus expression constructs for cytosolic Gamillus (top) and synaptophysin linked to Gamillus (bottom). D. Representative scatterplot showing FITC and BFP channels from flow cytometry analysis of iTF-Microglia and neuron cocultures. Two coculture configurations are displayed: a coculture of iTF-Microglia with a nuclear blue fluorescent protein marker and iNeurons with no engineering and a coculture of of iTF-Microglia with a nuclear blue fluorescent protein marker and iNeurons expressing synaptophysin linked to a green fluorescent protein, Gamillus. E. Gating strategy for isolating iTF-Microglia from coculture and a comparison of the FITC shift in iTF-Microglia cocultured with iNeurons (dark blue) and iTF-Microglia cocultured with iNeurons expressing synaptophysin linked to a green fluorescent protein, Gamillus (light blue). F. Representative micrographs iNeuron monocultures (left) and iTF-Microglia and iNeuron coculture (right) stained with the dead cell indicator, TO-PRO-3 (magenta), and nuclear stain, Hoechst 33342 (green). G. (Left) Percent live cells based on nuclear TO-PRO-3 intensity (N = 4 fields of view (FOV) for 5 wells, bars represent mean +/- standard deviation). (Right) Total cell count per field of view based on nuclear Hoechst segmentations (N = 4 fields of view for 5 wells, bars represent mean +/- standard deviation).

**Supplemental Figure 3.**
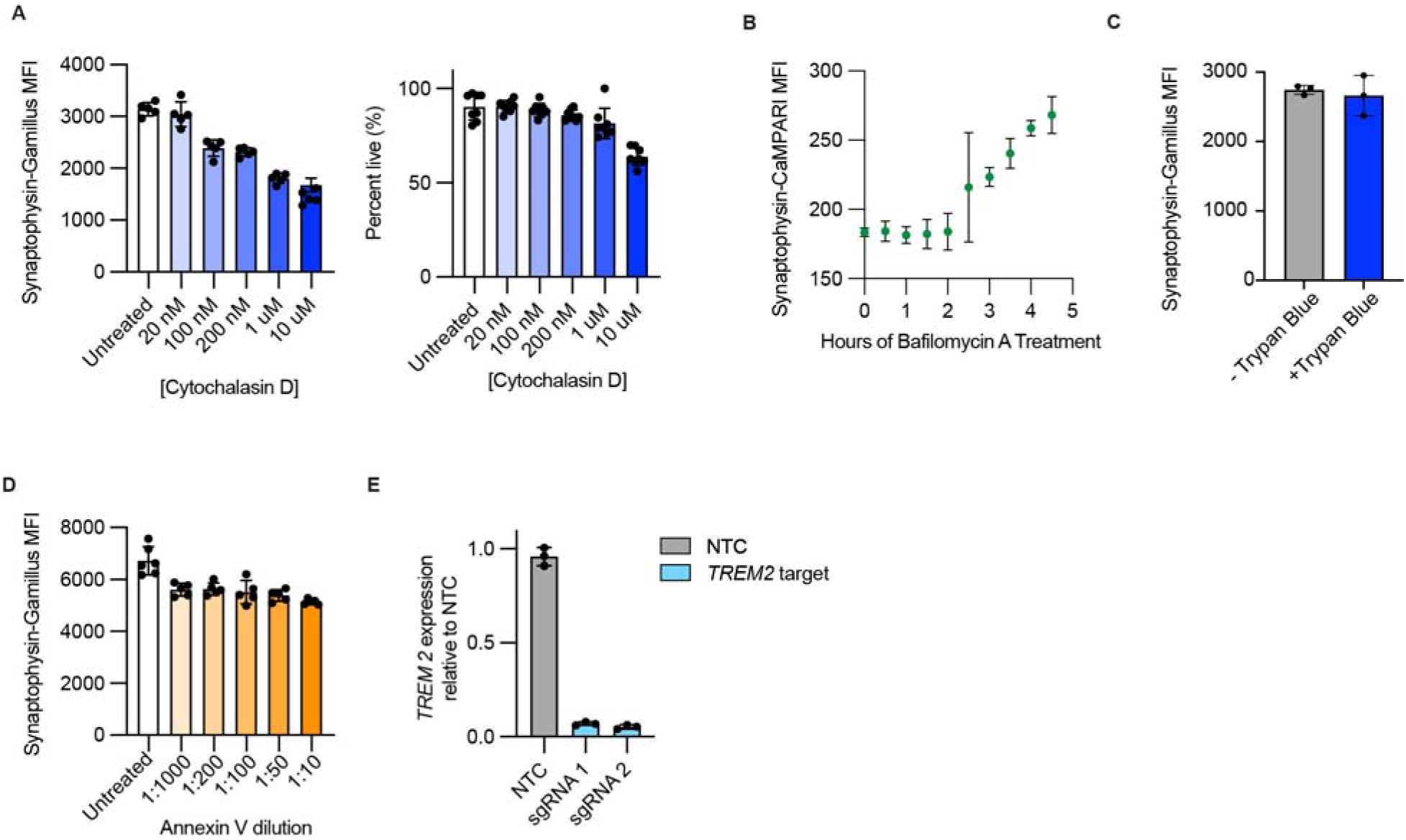
Characterization of synaptic pruning assay with pharmacological and genetic perturbations. A. (Left) Uptake of synaptic material by iTF-Microglia measured by flow cytometry after treating cocultures with increasing concentrations of Cytochalasin D (N = 5 wells, bars represent mean +/- standard deviation). (Right) Percent live cells based on nuclear TO-PRO-3 intensity with increasing concentrations of cytochalasin D (N = 4 fields of view from 2 wells, bars represent mean +/- standard deviation). B. Uptake of synaptic material by iTF-Microglia measured by flow cytometry after treating with Bafilomycin A for increasing incubation times. C. Uptake of synaptic material by iTF-Microglia measured by flow cytometry with and without Trypan Blue quenching. D. Uptake of synaptic material by iTF-Microglia measured by flow cytometry after treating cocultures with decreasing dilutions of annexin V (N = 5-6 wells, bars represent mean +/- standard deviation). E. *TREM2* mRNA expression in iTF-Microglia with sgRNAs targeting *TREM2* or non-targeting control (NTC) measured by qPCR.

**Supplemental Fig. 4:**
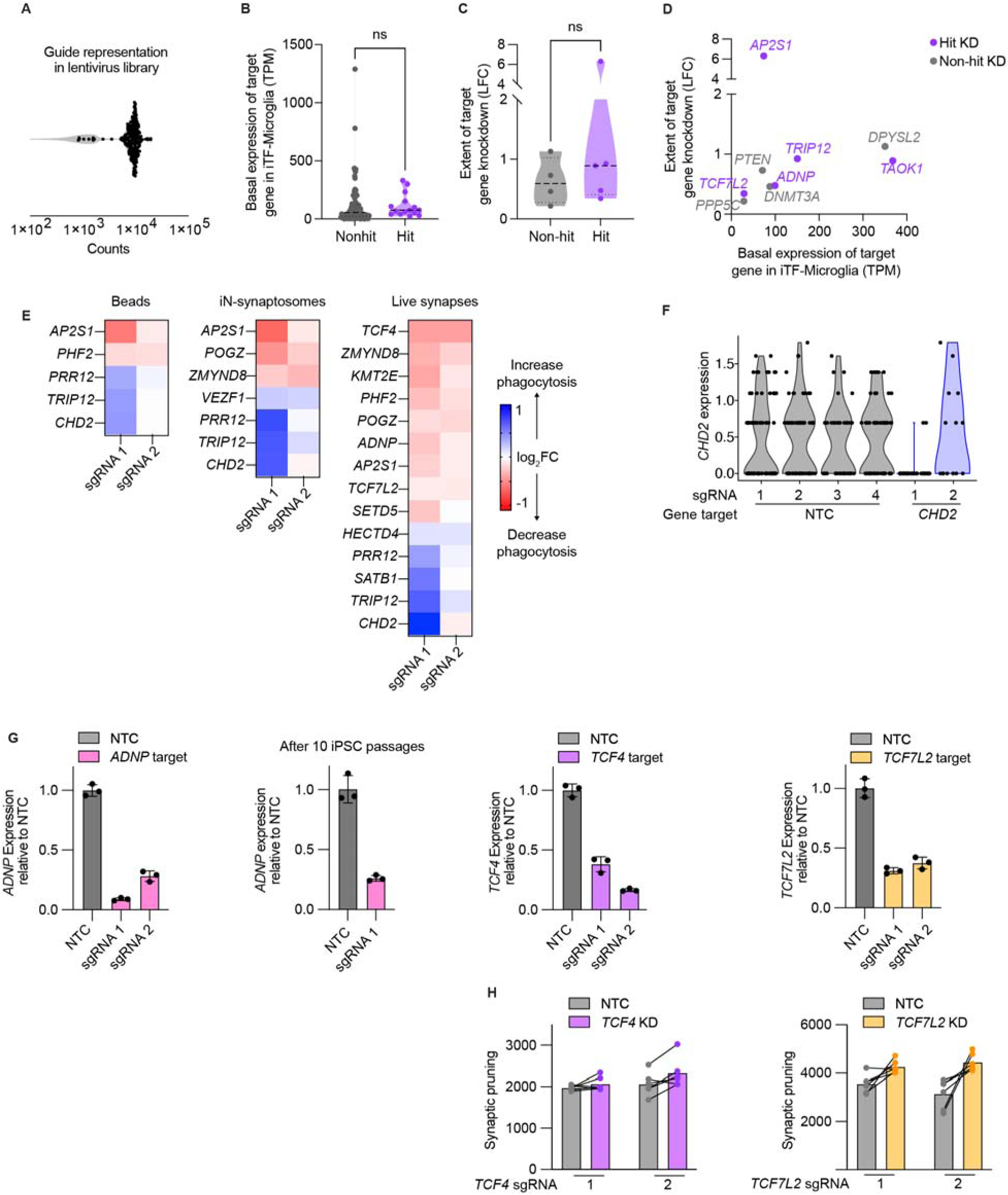
Robustness of hits across basal expression and sgRNAs. A. Distribution of sgRNA representation for CRISPRi-based ASD risk gene library. B. Comparison of iPSC-derived microglia (iTF-Microglia) basal expression of genes that emerged as hits or non-hits across all screens (two-tailed Student’s t-test). C. Comparison of knockdown (KD) efficacies based on single-cell transcriptomics dataset for genes that emerged as hits or non-hits across all screens (two-tailed Student’s t-test). D. Scatterplot aggregating iTF-Microglia basal expression, extent of KD, and hit classification for target genes. E. Heatmaps for each screen displaying the phagocytosis phenotype across both guides for genes that emerged as hits. F. *CHD2* expression level from the single-cell transcriptomics dataset for iTF-Microglia with non-targeting controls (NTCs) and iTF-Microglia with sgRNAs targeting *CHD2* (violin plot area represents the density curve). G. *ADNP, TCF4, TCF7L2* mRNA expression in iTF-Microglia with sgRNAs targeting those genes or NTC measured by qPCR. H. Synaptic pruning for iTF-Microglia with individual guides targeting *TCF4* and *TCF7L2*.

**Supplemental Fig. 5:**
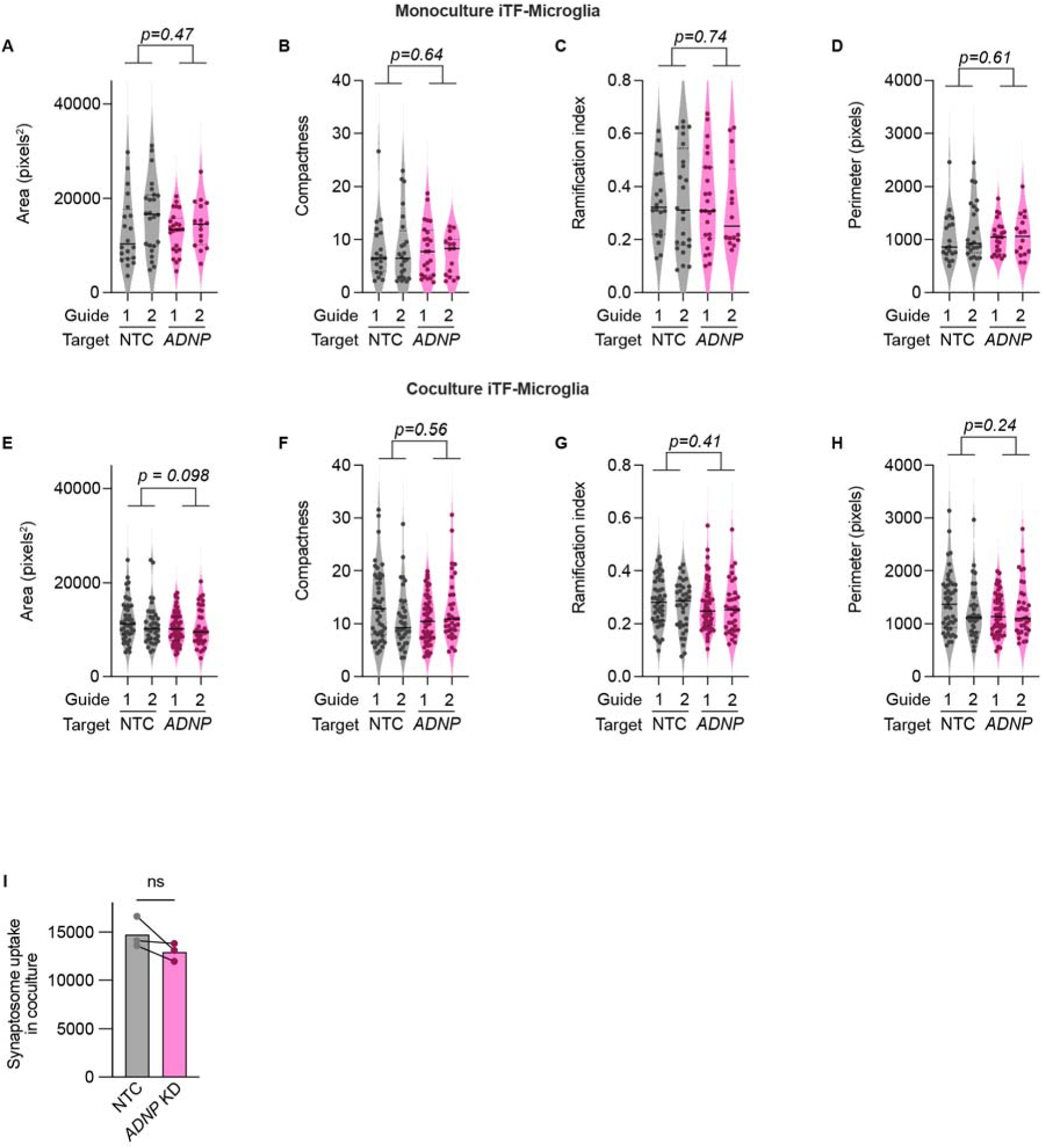
iPSC-derived microglia (iTF-Microglia) morphology and uptake in monoculture and coculture. A-D. Single cell morphology metrics for iTF-Microglia with *ADNP* KD or NTC in monoculture (N = 16-24 single cells, violin plot area represents the density curve, median and quartiles noted as bold and dashed lines respectively, two-way ANOVA). E-H. Single cell morphology metrics for iTF-Microglia with *ADNP* KD or NTC in coculture (N = 35-59 single cells, violin plot area represents the density curve, median and quartiles noted as bold and dashed lines respectively, significance based on NTC vs ADNP factor computed by two-way ANOVA). I. Uptake of iNeuron-derived synaptosomes by iTF-Microglia in coculture measured by flow cytometry (n = 3 wells, bars represent mean +/- standard deviation, paired t-test).

**Supplemental Fig. 6:**
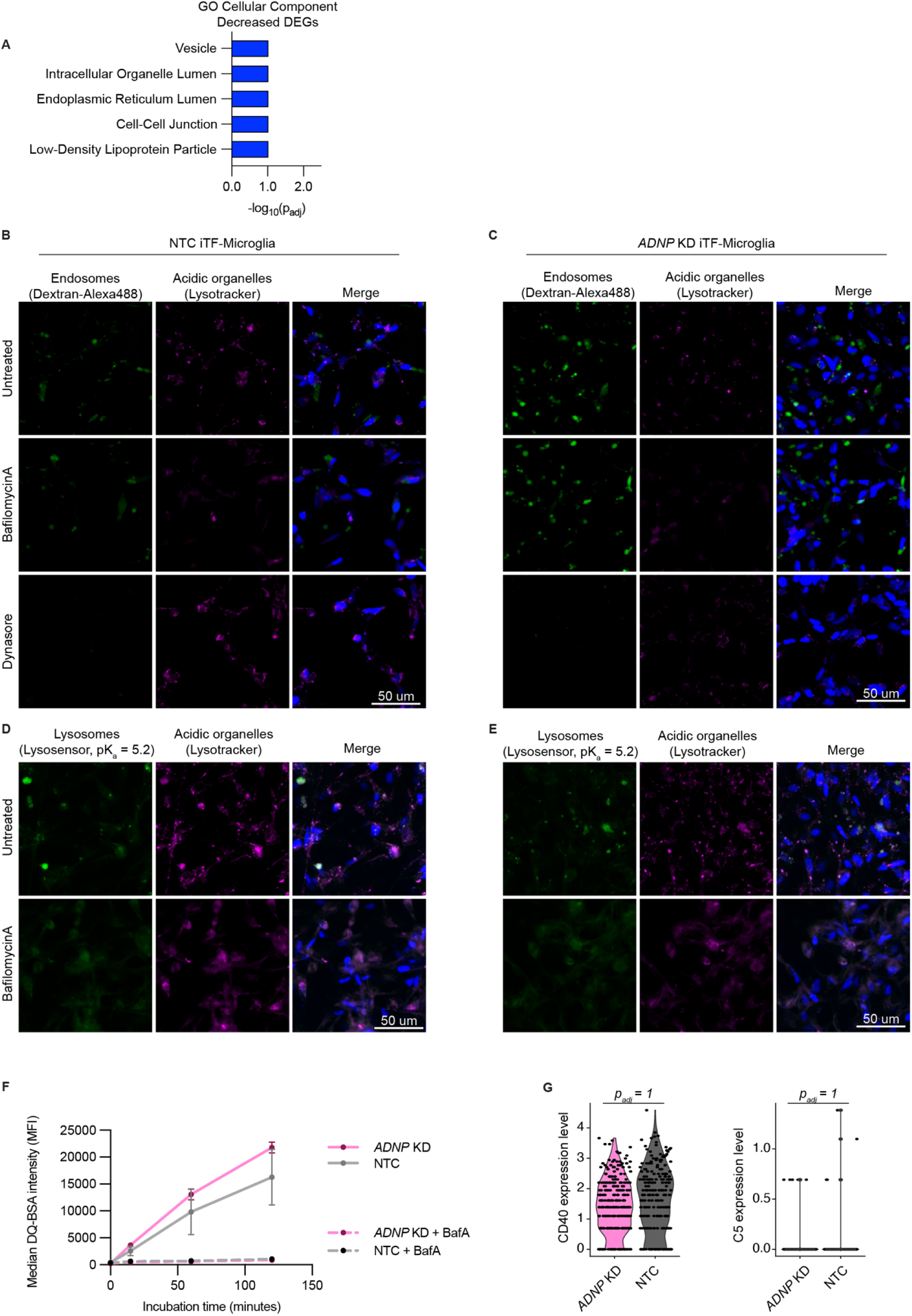
Endolysosomal characterization. A. GO Cellular Component enrichment analysis for downregulated differentially expressed genes for iTF-Microglia with *ADNP* knockdown (KD). B-C. Representative immunofluorescence micrographs of iTF-Microglia with non-targeting control (NTC) (A) or *ADNP* KD (B) incubated with Dextran-Alexa488 (green) and Lysotracker Deep Red (magenta) for 30 minutes with no treatment, BafilomycinA treatment, or Dynasore treatment. D-E. Representative immunofluorescence micrographs of iTF-Microglia with NTC (D) or *ADNP* KD (E) incubated with Lysosensor Green (green) and Lysotracker Deep Red (magenta) for 30 minutes with no treatment or BafilomycinA treatment. F. DQ-BSA fluorescence in iTF-Microglia with NTC (gray lines) or *ADNP* KD (pink lines) with increasing incubation times and BafilomycinA treatment (dashed lines) (N = 3 wells, bars represent mean +/- standard deviation, 2 way ANOVA. G. Expression level from the single-cell transcriptomics dataset for secreted proteins identified by the cytokine array to have increased abundance in iTF-Microglia with *ADNP* KD (violin plot area represents the density curve, two-tailed Student’s t-test).

**Supplemental Fig. 7:**
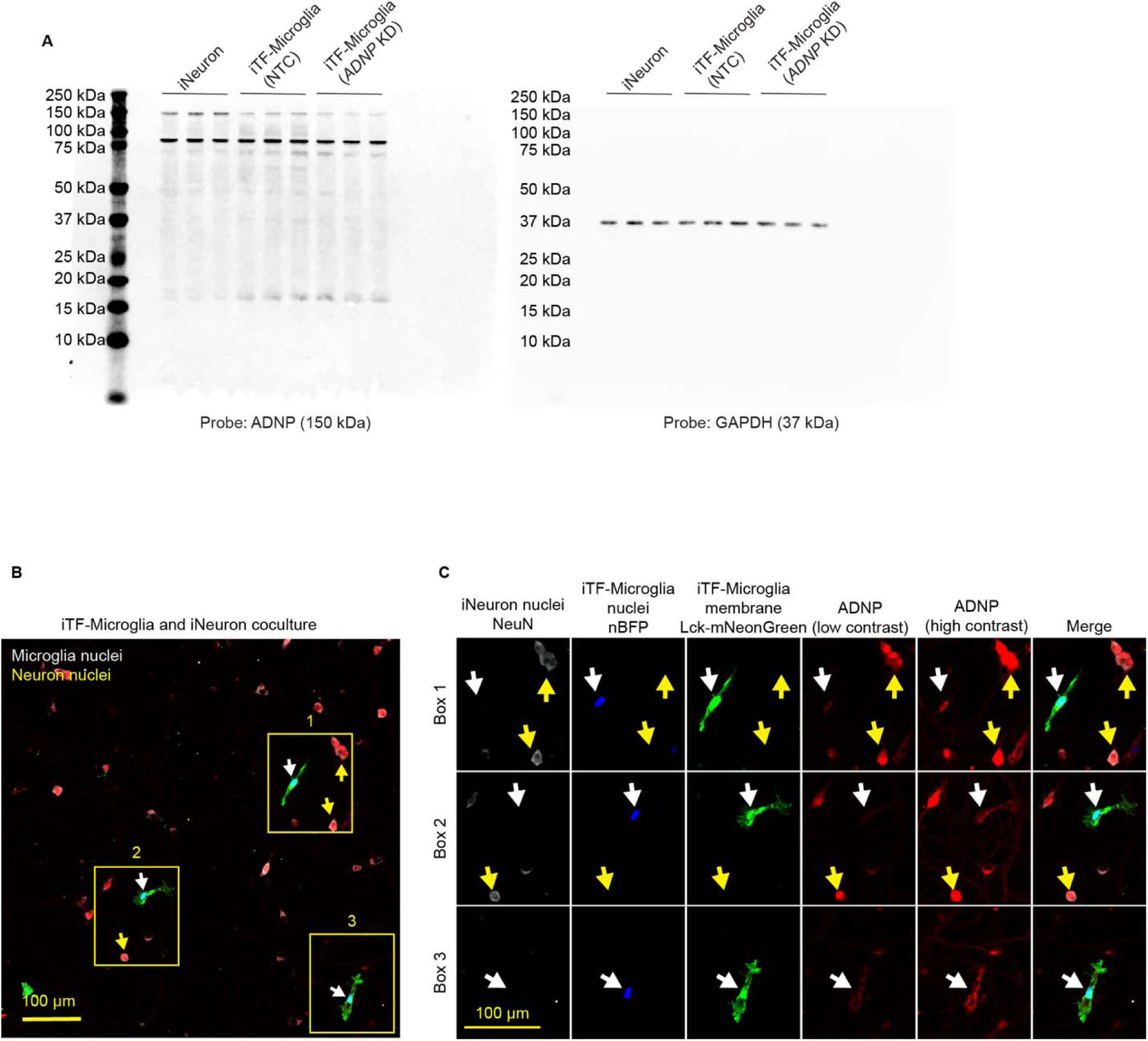
ADNP antibody and localization validation. A. Raw western blot images for ADNP stain (left) and GAPDH (right) for iPSC-derived neurons (iNeurons), iTF-Microglia with non-targeting control, and iTF-Microglia with *ADNP* knockdown (N = 3 replicates per cell type). B. Representative micrograph of iTF-Microglia expressing membrane marker, Lck-mNeonGreen (green) and nuclear BFP (blue), and iNeuron coculture stained with NeuN (gray) and ADNP (red). White arrows indicate microglia and yellow arrows indicate neurons that are shown magnified. C. Insets and individual channels visualizing three iTF-Microglia in the coculture and their ADNP expression.

**Supplementary Movie 1: Synaptophysin-Gamillus trafficking in iPSC-derived neurons (iNeurons).** iNeurons engineered to express synaptophysin-Gamillus were sparsely plated with unlabeled iNeurons. Individually engineered iNeurons show trafficking of synaptophysin-Gamillus puncta.

**Supplementary Movie 2: iPSC-derived microglia (iTF-Microglia) engulf synaptophysin-Gamillus puncta.** Labeled iTF-Microglia (gray) were cocultured with iPSC-derived neurons

(iNeurons) engineered to express synaptophysin-Gamillus (green). iTF-Microglia can engulf synaptophysin-Gamillus puncta.

**Supplementary Movie 3: iPSC-derived microglia (iTF-Microglia) traffick phagocytic cups containing synaptophysin-Gamillus.** Labeled iTF-Microglia (gray) were cocultured with iPSC-derived neurons (iNeurons) engineered to express synaptophysin-Gamillus (green). iTF-Microglia traffick phagocytic cups containing synaptophysin-Gamillus puncta toward their soma.

**Supplementary Table 1: sgRNA sequences targeting 102 ASD risk genes.**

**Supplementary Table 2: GO Cellular Component output for upregulated DEGs in Gandal 2022.**

**Supplementary Table 3: Raw cytokine array values for 30 secreted proteins.**

## Notes

### Summary of Updates

This version has additional experiments and analysis, and an improved logical flow

